# A Compact Regulatory Logic Governs Fibroblast Cell State Plasticity in Fibrosis

**DOI:** 10.64898/2026.10.09.757677

**Authors:** Ada M. Zhu, Yeting Li, Haiwei Zhang, Brindha Rathinasabapathi, Bastien Cimarosti, Rangarajan Nadadur, Vee Xu, Yu Huang, Yuqian An, Kirsten Auclair, Alicja Mahr, Maya U. Sheth, Clara Youngna Lee, Chen Wang, Tracy Yamawaki, Angelo Pelonero, Arun Padmanabhan, Sanjeev S. Ranade, Jesse M. Engreitz, Brandon Ason, Chi-Ming Li, Simon Jackson, Junedh M. Amrute, Pawel F. Przytycki, Michael Alexanian

## Abstract

Fibroblast activation drives organ failure across human diseases, yet its regulatory basis remains largely undefined. Here, we repressed 280 conserved disease-activated enhancers using CRISPR droplet sequencing (CROP-seq) in ∼550,000 fibroblasts, constructing a regulome of 15 gene programs governing the profibrotic state. Systematic TF perturbation converged on MEOX1 as a central regulator, and fibroblast-specific *Meox1* deletion in mice improved cardiac function, reduced fibrosis, and rewired profibrotic fibroblasts toward quiescence in heart failure. Effector gene perturbation in human cardiac fibroblasts revealed distinct regulators of contractility, migration, and inflammatory signaling, and these programs were conserved across profibrotic states in human lung, liver, and skin. Simultaneous repression of just six enhancers using a multiplexed dCas12a-KRAB system collapsed the profibrotic network, suppressed migration and contraction, and rewired fibroblast paracrine signaling to attenuate macrophage inflammation. These findings define the regulatory logic of fibroblast plasticity and establish a new framework for reprogramming profibrotic cell states in disease.

## INTRODUCTION

Tissue fibrosis is a hallmark of chronic disease, driving progressive organ failure in the heart, lungs, liver, kidneys, and skin and accounts for a substantial fraction of human morbidity and mortality worldwide^1^. Fibroblasts are central to this process, transitioning from homeostatic states into activated populations that deposit extracellular matrix (ECM), migrate, contract, proliferate, and signal to neighboring cells^2^. Although chronic fibroblast activation worsens dysfunction across fibrotic disorders and exacerbates many cancers, effective antifibrotic therapies remain largely unavailable^3,4^, and the few that exist slow disease progression without targeting the cell-intrinsic regulatory machinery that drives the profibrotic state. Importantly, fibrosis was long considered an irreversible endpoint, but growing evidence across organs now demonstrates that fibrotic remodeling is dynamic and that activated fibroblasts can revert to a quiescent-like phenotype when the underlying drivers are removed^5–8^. This reversibility suggests that the profibrotic state could be therapeutically reset by targeting the regulatory programs that sustain it.

Fibroblast activation encompasses multiple distinct biological functions, from ECM production to migration to inflammatory signaling, suggesting that the profibrotic state arises from a coordinated regulatory network of enhancers and transcription factors (TFs) rather than a single gene or pathway. Advances in single-cell and epigenomic technologies have mapped disease-associated cell states and regulatory elements at unprecedented resolution^9,10^, but which enhancers are functionally required, how they organize into gene programs, and whether the network can be disrupted through a limited set of regulatory nodes can only be resolved through systematic perturbation in a disease-relevant cellular context.

Here, we address these questions by constructing a comprehensive enhancer regulome of fibroblast activation in disease. We integrated epigenomic profiling of mouse and human hearts with CRISPRi-based repression of 280 conserved disease-activated enhancers coupled with single-cell transcriptomics (CROP-seq), resolving 15 gene programs that govern the major functions of activated fibroblasts. Motif enrichment within these enhancers nominated candidate TFs for functional testing, and systematic perturbation converged on MEOX1 as a central regulator of the profibrotic enhancer landscape. Fibroblast-specific deletion of *Meox1* in a mouse model of heart failure reduced fibrosis, improved cardiac function, and rewired fibroblasts from pathological activation toward the quiescent state. *Cis*-regulated effector genes controlled distinct components of the activated state, and the regulatory programs defined by the cardiac regulome were conserved across profibrotic fibroblast states in multiple organs and human fibrotic diseases.

Finally, we asked whether this complex regulatory network could be reduced to a minimal regulatory logic. Using a multiplexed dCas12a-KRAB system to simultaneously repress six enhancers, each governing a distinct biological function, we collapsed the profibrotic network and redirected fibroblasts toward quiescence, establishing that a complex disease-associated cellular state can be sustained and dismantled through a remarkably compact regulatory architecture.

## RESULTS

### Perturbation of disease-activated enhancers reveals the regulatory architecture of the profibrotic state

To identify the regulatory elements that govern fibroblast activation in disease, we integrated epigenomic data from mouse and human hearts^7,11,12^. Using chromatin accessibility from fibroblasts in healthy and diseased hearts, together with nascent transcription, BRD4 occupancy, and H3K27 acetylation profiling in mouse cardiac fibroblasts, we identified candidate enhancers preferentially active in disease-state fibroblasts (**Fig. 1A, Fig. S1A**). We then prioritized orthologous human elements that were accessible in cardiac fibroblasts from patients with acute myocardial infarction (AMI) or heart failure^11^, yielding 280 conserved disease-activated enhancers (**Fig. 1A, Fig. S1B**). To link these enhancers to target genes in human disease, we applied an enhancer-gene (E2G) predictive model^13^ to cardiac fibroblast multiome data from healthy donors and patients with acute myocardial infarction (AMI) or heart failure (HF) (**Fig. 1B**). Many E2G-linked genes were upregulated in disease, including the transcription factors (TFs) *MEOX1* and *ETV4* (**Fig. 1B**). To functionally validate these regulatory elements, we cloned the 280 enhancers into a reporter library and measured their transcriptional activity using a massively parallel reporter assay (MPRA)^14–16^ in mouse cardiac fibroblasts (**Fig. S1C-E)**. 150 of 280 enhancers (54%) showed increased reporter activity following TGF-β stimulation, a canonical profibrotic stimulus, confirming their preferential activation during fibroblast transition to a disease-associated state (**Fig. 1C, Fig. S1F,G**). TGF-β-induced enhancer activity was abolished by the BET bromodomain inhibitor JQ1^17^, which has potent antifibrotic effects in heart failure ^7,18–20^ (**Fig. S1H**). Motif analysis of TGF-β-responsive enhancers revealed enrichment for FOS-JUN/AP-1, ATF3, MEOX1, ERG, ETS1, ETV4, and RUNX1 motifs (**Fig. 1D**).

To determine the endogenous functions of these enhancers, we performed CRISPR droplet sequencing (CROP-seq)^21^ in mouse cardiac fibroblasts expressing dCas9 fused to the KRAB transcriptional repression domain (CRISPRi)^22^. We designed sgRNAs targeting within 500 bp of each enhancer’s midpoint, cloned them into a lentiviral CROP-seq library (including 20 non-targeting controls) covering all 280 enhancers, and transduced cells at low multiplicity of infection to enrich for cells carrying a single guide targeting a single enhancer. Cells were then treated with TGF-β and processed for single-cell 3’ gene expression profiling alongside sgRNA amplicon sequencing for guide assignment (**Fig. S2A**). After quality filtering, we obtained transcriptomic profiles from 549,767 single cells (**Fig. 1E**, **Fig. S2B–H**). To identify coordinated gene programs and the enhancers that drive them, we applied ALPINE^23^, a semi-supervised non-negative matrix factorization framework that extracts interpretable gene programs from multi-condition single-cell data (**Fig. S3A,B**). ALPINE identifies gene programs by decomposing single-cell expression data into sets of co-regulated genes, while simultaneously linking each program to the enhancer perturbations that alter it. The biological identity of each program was then assigned based on the top-ranked genes and pathway enrichment analysis. This resolved 15 gene programs organized into ten biological categories that collectively define the functional landscape of activated fibroblasts (**Fig. 1F–G**, **Fig. S3B-D**). These included contractile myofibroblast programs marked by *Tagln*, *Myl9*, and TGF-β autocrine signaling; ECM deposition and maturation (*Ptn*, *Bgn*); migration and paracrine signaling (*Svil*, *Has2*, *Cxcl12*); myofibroblast proliferation programs including a *Postn*-positive activated population; ECM sensing and integrin-mediated turnover; chromatin remodeling and stress adaptation; metabolic activation; and a quiescent immunomodulatory state (*Cdkn1a*, *Klf2*, *Ccl2*). Each program was linked to the specific enhancers whose CRISPRi-mediated repression altered its activity, providing both a transcriptional and a regulatory map of disease-associated fibroblast cell states (**Fig. S3E**). To determine whether the genes driving these programs are modulated in cardiac disease, we scored their activity in human and mouse datasets^7,11^. In human cardiac fibroblasts, most profibrotic programs were strongly induced in AMI, consistent with the broad activation of fibroblasts following ischemic injury. The migratory-paracrine (P7) and biosynthetic (P2) programs were instead preferentially active in chronic heart failure, suggesting distinct regulatory dynamics between acute and chronic disease. The cytoprotective (P11) and quiescence (P1) programs showed the opposite pattern, with highest activity in healthy donors, indicating that these homeostatic states are lost upon injury (**Fig. 1H, left**). In mice subjected to pressure-overload heart failure by transverse aortic constriction (TAC), the profibrotic programs, including contractile (P5, P6, P10), ECM deposition (P9), myofibroblast proliferation (P13, P14), and ECM sensing (P15), were induced with disease, suppressed by the BET bromodomain inhibitor JQ1, and re-emerged after treatment withdrawal, tracking with the extent of cardiac fibrosis^7^ (**Fig. 1H, right**). Conversely, the quiescence program (P1) was enriched in sham hearts, lost in disease, restored by anti-fibrotic treatment, and suppressed again after withdrawal, highlighting fibroblast activation and quiescence as opposing, dynamically reversible transcriptional states (**Fig. 1H**).

**Figure 1.**
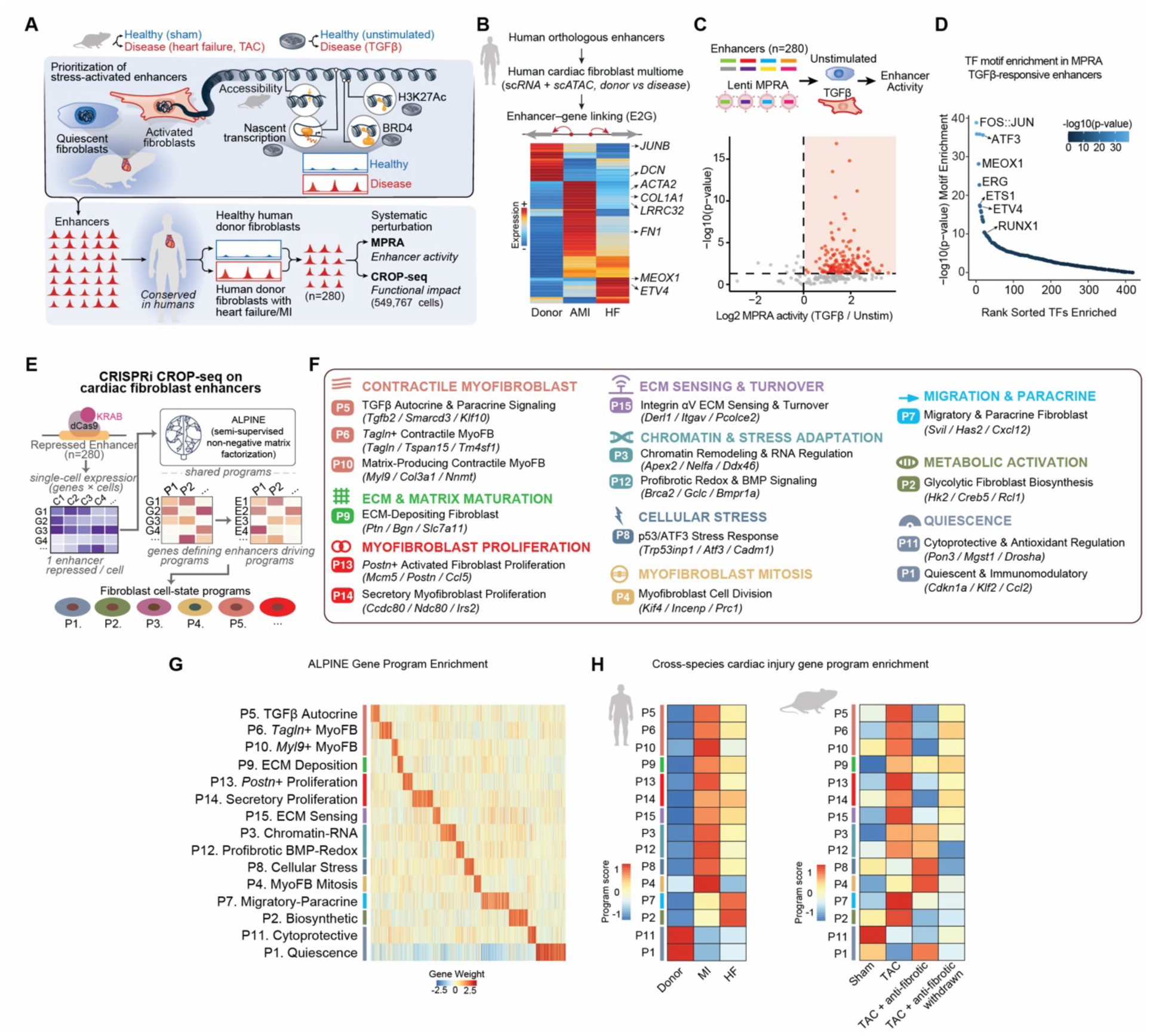
Functional interrogation of cardiac fibroblast enhancers defines cell-state gene programs. **A.** Integrative approach for prioritization of stress-activated enhancers in cardiac fibroblasts from *in vivo* (Sham vs. TAC) and *in vitro* (Unstim vs. TGFβ) comparisons. Conserved human orthologous enhancers (n=280) were tested by MPRA and CRISPRi CROP-seq (549,767 cells) in mouse cardiac fibroblasts. **B.** Expression of enhancer-linked genes across non-failing (Donor), acute myocardial infarction (AMI), and heart failure (HF) human cardiac fibroblasts, derived from multiome (scRNA + scATAC) with enhancer–gene (E2G) linking. **C**. Lentiviral MPRA activity of 280 enhancers in unstimulated and TGFβ-treated mouse cardiac fibroblasts. Log_2_ fold change (TGFβ/Unstim) versus −log_10_(p-value). **D.** TF motif enrichment in TGFβ-responsive MPRA enhancers. Top enriched TFs highlighted; inset shows −log_10_(p-value). **E.** CRISPRi CROP-seq schematic. dCas9-KRAB repression of individual enhancers (1 per cell) coupled with single-cell RNA-seq and ALPINE (semi-supervised non-negative matrix factorization) to define fibroblast cell-state programs. **F.** Fifteen fibroblast cell-state programs (P1–P15) identified by ALPINE, organized by functional category with representative genes. **G.** Heatmap of ALPINE gene weights across the 15 fibroblast programs. **H**. Cross-species cardiac injury gene program enrichment in human cardiac fibroblasts (Donor, MI, HF; left) and mouse cardiac fibroblasts (Sham, TAC, TAC + anti-fibrotic (JQ1), TAC + anti-fibrotic (JQ1) and then JQ1 withdrawal; right).

Together, these results define a regulome of disease-activated enhancers that govern distinct fibroblast gene programs, validated across species and disease models, and define the regulatory architecture through which fibroblasts transition from quiescence to pathological activation in disease.

### A transcription factor hierarchy governs the profibrotic regulatory network

We next asked which enhancers are most critical for maintaining the disease-associated fibroblast state. We trained logistic regression classifiers on single-cell RNA-seq data from healthy and diseased mouse and human hearts to score cells along a healthy-to-disease axis, then used the learned feature weights to quantify how repression of each enhancer in the CROP-seq dataset shifted cells toward a healthier transcriptional state (**Fig. 2A, Fig. S4A–C**). Enhancers whose repression concordantly shifted fibroblasts toward a healthy transcriptional state in both the mouse and human classifiers were defined as profibrotic enhancers (**Fig. 2B**, **Fig. S4D**). To link the mouse profibrotic enhancers interrogated by CROP-seq to their target genes, we generated H3K27ac ChIP-seq and ATAC-seq profiles in mouse cardiac fibroblasts and built an E2G model^13^. Predicted targets included established fibrotic genes such as *Postn*, *Col1a1*, and *Serpine1*, as well as less characterized genes including *Nrep*, *Adam12, Lrrc32*, and *Edn1*. The profibrotic enhancers were among the most strongly TGF-β-responsive elements in the MPRA, and their linked genes were induced in human heart failure and AMI relative to healthy donors, confirming that disease-driving regulatory activity, intrinsic transcriptional potential, and target gene expression converge at these loci (**Fig. 2C, Fig. S4E**). We then asked how the profibrotic enhancers were distributed across the fibroblast gene programs. The ECM deposition program (P9) was the most enriched, with over half of its enhancers classified as profibrotic, followed by the profibrotic redox and BMP signaling (P12) and ECM sensing and turnover (P15) programs (**Fig. 2D**).

We next sought to determine which TFs regulate this profibrotic enhancer network. Motif enrichment analysis ranked MEOX1 as the most enriched TF, followed by FOS::JUN, MAFG, BACH1, and PRDM4 (**Fig. 2E**). Six of the top-ranked factors, MEOX1, c-FOS, c-JUN, RUNX1, ETS1, and ETV4, were also among the most enriched motifs in the TGF-β-responsive MPRA enhancers (**Fig. 1D**), and all were expressed in cardiac fibroblasts in both mouse and human heart failure (**Fig. S5A,B**). To test the role of each TF in governing the profibrotic regulatory network, we individually knocked down each of the six factors using CRISPRi in cardiac fibroblasts, stimulated cells with TGF-β, and profiled transcriptional and chromatin changes by bulk RNA-seq and ATAC-seq (**Fig. 2F, S5C-G**). MEOX1 depletion had the broadest chromatin effect, reducing accessibility at approximately 22% of profibrotic enhancers, followed by RUNX1 (**Fig. 2G, top**). Furthermore, through E2G mapping, we found that MEOX1, ETV4, and RUNX1 repression abrogated a substantial fraction of profibrotic target gene expression (**Fig. 2G, bottom**). Notably, ETV4 loss had minimal effect on enhancer accessibility despite strongly altering target gene expression, suggesting that ETV4 regulates transcription through a mechanism distinct from the broader chromatin remodeling exerted by MEOX1 and RUNX1. We next examined which fibroblast gene programs were most sensitive to each TF perturbation at the chromatin level. MEOX1 loss produced the broadest reduction in accessibility at enhancer-linked target genes, with the strongest effect on the ECM deposition (P9) and the contractile myofibroblast programs (P5, P6, P10). RUNX1 showed a more selective pattern, primarily affecting accessibility at genes within the migratory-paracrine program (P7). c-FOS loss also broadly impacted several programs, with a notably strong effect on the metabolic activation program (P2) (**Fig. 2H**). To illustrate this regulatory architecture at individual loci, we examined three representative profibrotic enhancer-gene pairs: *Adam12*, *Nrep*, and *Serpine1*.

**Figure 2.**
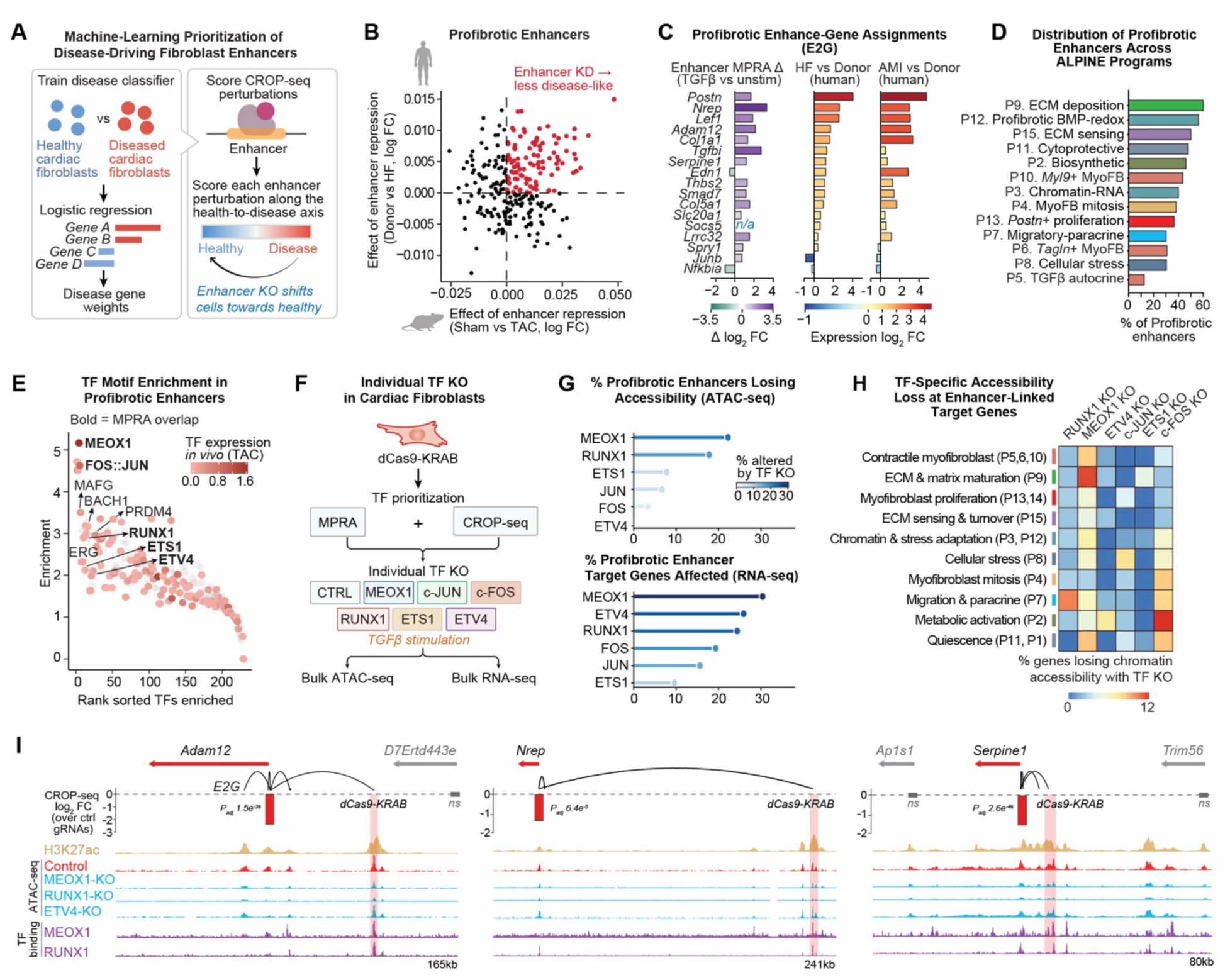
Disease-driving enhancers converge on a defined set of transcription factor regulators. **A.** Schematic of disease classifier approach. Logistic regression trained on healthy vs. disease cardiac fibroblasts assigns disease gene weights; CRISPRi CROP-seq perturbations scored by shift along the healthy-to-disease axis. **B**. Disease score of each enhancer in mouse (x-axis, Sham vs. TAC) and human (y-axis, Donor vs. HF) classification frameworks. Profibrotic enhancers (red) shift fibroblasts toward a healthier state upon CRISPRi repression in both species. **C**. Profibrotic enhancer–gene assignments (E2G) showing enhancer MPRA activity change (TGFβ vs. Unstim), and expression of linked genes in human HF vs. Donor, and AMI vs. Donor in cardiac fibroblasts. **D.** Distribution of profibrotic enhancers across ALPINE programs. **E.** TF motif enrichment in profibrotic enhancers. Bold indicates MPRA overlap. Top enriched TFs highlighted. **F.** Schematic for individual TF knockdown in cardiac fibroblasts. dCas9-KRAB targeting of TFs prioritized by MPRA and CROP-seq (MEOX1, c-JUN, c-FOS, RUNX1, ETS1, ETV4) followed by TGFβ stimulation, bulk ATAC-seq and bulk RNA-seq. **G.** Percentage of profibrotic enhancers losing chromatin accessibility (ATAC-seq) with each TF knockdown (top) and percentage of profibrotic enhancer linked genes affected (bottom). **H.** TF-specific accessibility loss at enhancer-linked target genes across ALPINE programs. **I.** Representative loci showing CROP-seq log_2_ FC (over control gRNAs), H3K27ac, bulk ATAC-seq in control and TF knockdown conditions, and TF binding (MEOX1, RUNX1) at the *Adam12*, *Nrep* and *Serpine1* loci. E2G links and dCas9-KRAB target sites indicated.

CRISPRi-mediated repression of each enhancer significantly downregulated its linked gene in the CROP-seq data, confirming bona fide *cis*-regulatory relationships supported by E2G assignments and H3K27ac marking (**Fig. 2I, top**). MEOX1 ChIP-seq and RUNX1 CUT&RUN confirmed direct binding of both TFs at these enhancers (**Fig. 2I**, **bottom**, **Fig. S5H,I**). Loss of MEOX1 or RUNX1 reduced chromatin accessibility at both the enhancer and the promoter of each target gene, consistent with a model in which these TFs maintain an accessible chromatin state across the full *cis*-regulatory unit (**Fig. S5J**). ETV4 repression by contrast had little effect on enhancer accessibility but reduced accessibility at the *Adam12* promoter, consistent with regulation of target gene expression without widespread chromatin remodeling (**Fig. 2I**). Together, these findings reveal a hierarchical regulatory architecture of fibroblast activation, in which a profibrotic enhancer network is coordinated by distinct TFs, with MEOX1 acting as a dominant upstream regulator of the disease-associated state.

### *Meox1* is required for fibroblast activation *in vivo*

MEOX1 emerged from the CROP-seq regulome as the broadest regulator of profibrotic enhancer activity, affecting chromatin accessibility, target gene expression, and multiple fibroblast programs. Prior studies have shown that, in acute and chronic human heart failure, *MEOX1* is expressed in profibrotic *POSTN*^+^ fibroblasts^11,12^ (**Fig. S6A**). Our previous work had shown that *Meox1* is induced in cardiac fibroblasts during pressure overload, suppressed by BET inhibition, and required for fibroblast activation *in vitro*^7,24^. However, whether *Meox1* is required for fibrosis and cardiac dysfunction *in viv*o, and which fibroblast gene programs it controls in the diseased heart, remained unknown. We generated a mouse line harboring a conditional allele in which loxP sites flank the entire *Meox1* coding region (**Fig. S6B**). To validate the allele, we first crossed it with a β-actin-Cre. Consistent with the established role of *Meox1* in somitogenesis^25,26^, global knockout mice showed loss of *Meox1* expression, sub-Mendelian survival, and skeletal deformities in surviving animals (**Fig. S6C-E**). To specifically delete *Meox1* in disease-activated cardiac fibroblasts, we crossed the conditional allele with a tamoxifen-inducible Cre driven by the promoter of *Postn*^27^, and subjected mice to pressure overload by TAC (**Fig. 3A–B, Fig. S6F**). Activated fibroblast-specific *Meox1* deletion improved cardiac function, as measured by global longitudinal strain and left-ventricular ejection fraction, and reduced cardiac fibrosis at six weeks after TAC (**Fig. 3C–E**). To investigate the transcriptional mechanism of this cardioprotective phenotype, we performed single-cell RNA-seq on the non-cardiomyocyte fraction at six weeks post-TAC (**Fig. 3F, Fig. S6G,H)**. *Meox1* expression was enriched in fibroblasts and notably absent from cardiomyocytes as measured by RNA expression and RNAscope (**Fig. 3F–H, Fig. S6I-K**). Consistent with this, fibroblasts showed the greatest transcriptional response to *Meox1* deletion, with the largest number of differentially expressed genes across cell types, followed by macrophages (**Fig. 3I**). Gene ontology analysis of the genes downregulated in *Meox1* knockout fibroblasts revealed enrichment for pathways central to pathological activation, including response to ER stress, regulation of TGF-β signaling, ECM organization, and supramolecular fiber organization (**Fig. 3J**). To dissect the gene programs controlled by *Meox1 in vivo*, we applied consensus non-negative matrix factorization (cNMF)^28^ to decompose the fibroblast transcriptomes from control and *Meox1* conditional-knockout hearts into coordinately expressed gene programs (**Fig. 3K**). This identified seven programs (GP1–GP7), each defined by a distinct biological signature (**Fig. 3L**). GP2, the dominant profibrotic module, contained *Meox1* alongside *Ctgf*, *Postn*, and the less characterized gene *Mical2*. A second ECM program, GP7, featured the matricellular genes *Fmod*, *Comp, Cilp2* and *Thbs1*. GP6 was characterized by genes involved in cell-surface interactions including *Vim* and *Dkk3*. In contrast, GP3 was defined by immediate-early transcription factors associated with quiescence (*Junb*, *Fos*), GP4 represented a resting fibroblast identity marked by *Pi16* and the stromal marker *Cd248*, and GP5 was enriched for secreted homeostatic factors such as *Svep1*, *Sfrp1*, and *Fbln1*. Comparison of program usage between control and *Meox1*-deleted fibroblasts revealed a profound remodeling of the *in vivo* transcriptional landscape (**Fig. 3M**). The profibrotic ECM program GP2 was the most strongly depleted upon *Meox1* loss, along with GP6 and GP7. Conversely, GP3, GP4, and GP5 were all significantly gained, including the quiescence-associated programs GP3 and GP4. *Meox1* deletion therefore does not simply attenuate profibrotic gene expression, it actively redirects fibroblasts from a pathological ECM-producing state toward quiescence and a homeostatic identity. To distinguish direct transcriptional targets of MEOX1 from changes secondary to improved cardiac function, we integrated the *in vivo* gene programs with MEOX1 ChIP-seq binding data^7^ (**Fig. 3N**). GP2 showed the strongest enrichment for MEOX1-bound genes and GP7 was also enriched, supporting direct MEOX1 regulation of both ECM programs. Notably, GP6, though depleted upon *Meox1* deletion, was not enriched for MEOX1 binding, suggesting that its loss is a secondary rather than direct consequence of *Meox1* loss. The quiescence and homeostatic programs gained upon *Meox1* deletion (GP3 and GP4) were similarly depleted for MEOX1 binding, indicating that the reactivation of these programs is an indirect consequence of dismantling the MEOX1-driven profibrotic state. To determine how the *in vivo* programs correspond to the enhancer-defined programs identified by CROP-seq, we computed the similarity between the seven *in vivo* gene programs and the 15 *in vitro* CROP-seq programs (**Fig. 3O**). The MEOX1-regulated profibrotic module GP2 mapped directly onto the CROP-seq ECM deposition program P9, the program most enriched for profibrotic enhancers. GP7 also corresponded to P9 but additionally aligned with the secretory proliferation (P14) and migratory-paracrine (P7) programs, revealing that this *in vivo* module spans multiple dimensions of fibroblast pathology that were resolved as distinct programs *in vitro*. Remarkably, the quiescence programs GP3 and GP4, which emerged *in vivo* only upon *Meox1* deletion, mapped onto P1, the quiescence program identified independently through enhancer perturbation in the CROP-seq screen. Finally, we asked which of the genes suppressed by *Meox1* deletion *in vivo* are located at loci where the CROP-seq screen identified *cis*-regulatory enhancers bound by MEOX1. (**Fig. 3P**). This set included established profibrotic genes such as *Serpine1*, *Ctgf*, *Sertad4* and *Col5a1*, alongside *Adam12*, *Mical2*, *Aebp1*, *Nrep*, *Tgfbi*, and *Rbpj*.

**Figure 3.**
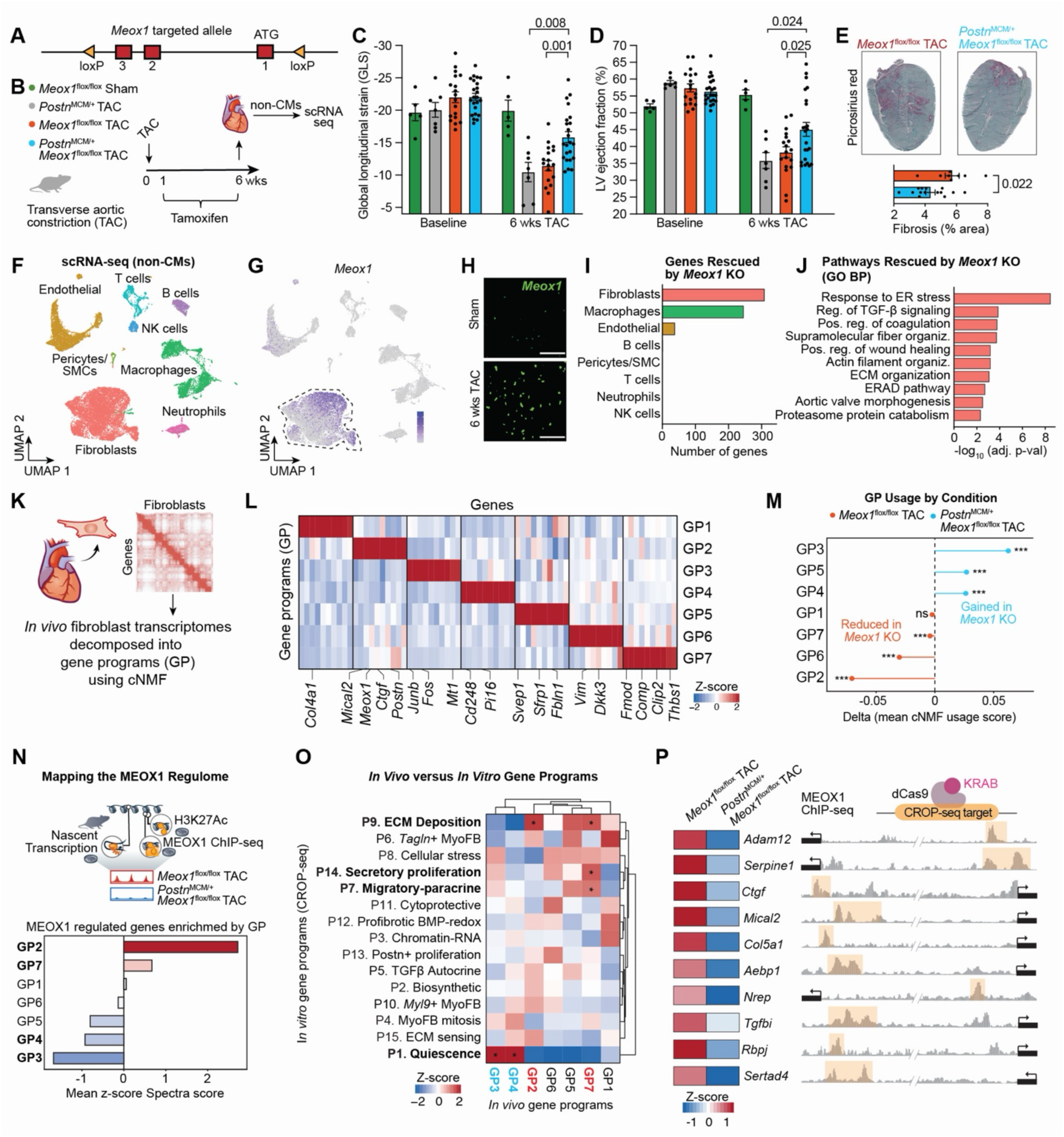
*Meox1* is required for cardiac profibrotic gene program activation *in vivo*. **A.** Schematic of the Meox1 targeted allele with loxP sites flanking exons 1–3. **B.** Experimental design. *Meox1*^flox/flox^ Sham, *Postn*^MCM/+^ TAC, *Meox1*^flox/flox^ TAC, *Meox1*^flox/flox^ ; *Postn*^MCM/+^ TAC mice subjected to transverse aortic constriction (TAC) with tamoxifen administration; non-CMs collected at 6 weeks for scRNA-seq. **C,D.** Global longitudinal strain (GLS) (C) and LV ejection fraction (D) at baseline and 6 weeks post-TAC. n=5 (*Meox1*^flox/flox^ Sham), n=7 (*Postn*^MCM^ TAC), n=18 (*Meox1*^flox/flox^ TAC), n=23 (*Meox1*^flox/flox^ ; *Postn*^MCM/+^ TAC). One-way ANOVA with post-hoc multiple comparisons; mean ± SEM. **E**. Picrosirius red staining and quantification of fibrosis (% area) in *Meox1*^flox/flox^ TAC and *Postn*^MCM/+^; *Meox1*^flox/flox^ TAC hearts. Unpaired t-test with Welch’s correction; mean ± SEM. **F.** UMAP of non-cardiomyocyte cells from scRNA-seq colored by cell type. **G.** UMAP of *Meox1* expression. **H.** *Meox1* RNAscope in Sham and 6-week TAC hearts. Scale bar = 50 μm. **I.** Number of genes rescued by *Meox1* KO across cell populations. **J.** GO biological process pathways rescued by *Meox1* KO in fibroblasts. **K.** Schematic of *in vivo* fibroblast transcriptome decomposition into gene programs using cNMF. **L.** Seven gene programs (GP1–GP7) with heatmap of representative genes. **M.** Gene program usage by condition. Delta mean cNMF usage score between *Meox1*^flox/flox^ TAC and *Postn*^MCM/+^; *Meox1*^flox/flox^ TAC. **N.** Mapping the MEOX1 regulome by integrating nascent transcription, H3K27Ac and MEOX1 ChIP-seq. Bar plot shows mean z-score Spectra enrichment of MEOX1-regulated genes across gene programs. **O.** Comparison of *in vivo* (cNMF) and i*n vitro* (CROP-seq ALPINE) gene programs. Heatmap of z-scores; asterisks denote significant overlaps. **P.** Expression z-scores of selected MEOX1-regulated genes in *Meox1*^flox/flox^ TAC and *Postn*^MCM/+^; *Meox1*^flox/flox^ TAC fibroblasts from scRNA-seq (left). Right: genome browser tracks showing MEOX1 ChIP-seq binding at each locus, with the CROP-seq targeted enhancer highlighted.

Thus, MEOX1 directly maintains the profibrotic network through defined *cis*-regulatory elements, and in its absence, fibroblasts do not adopt a novel transcriptional state but instead default to the quiescent program that the CROP-seq regulome identified as the transcriptional opposite of profibrotic activation. This convergence between fibroblast-specific deletion *in vivo* and systematic enhancer perturbation *in vitro* establishes MEOX1 as a pivot between the profibrotic and quiescent fibroblast states.

### *Cis*-regulated effector genes control distinct components of the profibrotic state

Having established the enhancer regulome, the TF hierarchy that governs it, and the *in vivo* role of MEOX1, we next investigated the mechanisms through which these enhancers regulate fibroblast function. For each of the 280 enhancers repressed by CRISPRi in the CROP-seq screen, we examined the expression of all genes within a 500-kb window. Affected genes were concentrated near the targeted elements, and downregulated genes substantially outnumbered upregulated ones, consistent with these regulatory elements functioning as bona fide transcriptional enhancers that activate nearby genes in *cis* (**Fig. 4A**). To identify the *cis*-regulated effector genes that mediate the distinct fibroblast programs revealed by the enhancer-based CROP-seq screen, we selected ten genes whose expression was significantly reduced upon repression of their linked enhancer and individually depleted each by siRNA in primary human cardiac fibroblasts, followed by bulk RNA-seq (**Fig. 4B, Fig. S7A-D**). Each of the ten depletions produced a distinct transcriptional signature, but four genes emerged as key regulators of distinct aspects of fibroblast activation (**Fig. 4C–D**). Depletion of *SERPINE1*, encoding the plasminogen activator inhibitor PAI-1, a well-established driver of ECM accumulation across fibrotic organs^29,30^, broadly reduced ECM gene expression. *ADAM12* loss suppressed both ECM and contractile programs, suggesting a cell-autonomous profibrotic function for a metalloproteinase that has not been studied in this context in cardiac fibroblasts. *MICAL2* depletion selectively reduced contractile and migratory signatures, and *AEBP1* (encoding ACLP) loss abolished an inflammatory transcriptional program. These results reveal that individual *cis*-regulated effector genes control distinct components of the activated fibroblast state.

To validate these predictions functionally, we performed collagen-gel contraction assays in primary human cardiac fibroblasts. Depletion of either *ADAM12* or *MICAL2* significantly reduced contraction (**Fig. 4E**). In trans-well migration assays, *MICAL2* depletion markedly impaired fibroblast migration (**Fig. 4F**), whereas *ADAM12* depletion had no effect (**Fig. S7E**), consistent with the selective migratory signature observed only upon *MICAL2* loss in the transcriptional analysis (**Fig. 4C,D**). MICAL2, a flavoprotein oxidoreductase that regulates actin dynamics, had no established role in cardiac fibrosis, making its control of fibroblast contractility and migration a previously unrecognized function. Because *AEBP1* depletion suppressed an inflammatory transcriptional signature rather than an ECM or contractile program, we asked whether this inflammatory response has paracrine consequences. Conditioned medium from *AEBP1*-depleted human primary cardiac fibroblasts was transferred to mouse bone marrow-derived macrophages, which showed reduced expression of the proinflammatory genes *Il6*, *Csf3*, and *Cxcl3* compared to macrophages exposed to control-conditioned medium, suggesting that *AEBP1*-dependent fibroblast signaling modulates macrophage inflammatory activation (**Fig. 4G**). Notably, *Adam12*, *Mical2*, and *Aebp1* were each repressed by *Meox1* deletion *in vivo* and their linked enhancers were bound by MEOX1 in the CROP-seq screen (**Fig. 3P**), placing these functionally specialized effector genes within the MEOX1-regulated profibrotic network. To determine whether the *cis*-regulatory mechanisms identified in the CROP-seq screen are conserved in humans, we performed bulk ATAC-seq in human cardiac fibroblasts and confirmed accessible, active regulatory elements at the orthologous loci near *ADAM12*, *MICAL2*, and *AEBP1* (**Fig. 4H**).

### The profibrotic regulatory architecture is conserved across organs and human fibrotic diseases

Fibroblast activation drives tissue remodeling and organ failure not only in the heart but across a wide range of human diseases. We therefore asked whether the regulatory programs and effector genes defined by the cardiac CROP-seq regulome are conserved across organs. To address this, we analyzed a cross-organ fibroblast atlas encompassing 220,179 fibroblasts across 16 tissues, including lung, liver, skin, intestine, and skeletal muscle, spanning 28 disease and fibrotic conditions and 13 recurrent cell states^31^ (**Fig. 4I,J**). Among these, the *LRRC15*-positive myofibroblast population represents a universal disease-emergent profibrotic state^32^, while *PI16*-positive and *COL15A1*-positive fibroblasts mark conserved homeostatic populations (**Fig. 4J**). The majority of the cis-regulated effector genes characterized in human cardiac fibroblasts, including *ADAM12*, *MICAL2*, *BGN*, and *COL5A1*, were preferentially expressed in the *LRRC15*-positive profibrotic state across organs, while homeostatic *PI16*-positive and *COL15A1*-positive fibroblasts showed minimal expression of these genes (**Fig. 4K**). The effector genes through which disease-activated enhancers control cardiac fibroblast behavior are therefore not restricted to the heart but are defining features of a universal profibrotic fibroblast identity. We next asked whether the profibrotic gene programs governed by disease-activated enhancers in the heart are also conserved across organs. P9, the ECM deposition program containing the largest share of profibrotic enhancers, showed the highest activity in the *LRRC15*-positive profibrotic population across all fibroblast states in both mouse and human (**Fig. 4L–M**). The transcriptional architecture defined by the cardiac fibroblast enhancer regulome is therefore a conserved feature of the profibrotic fibroblast state across species and tissues. To test whether these programs are dynamically regulated in human fibrotic disease, we scored their activity across disease stages in lung, liver, and skin (**Fig. 4N, Fig. S8A-C**). In the lung, P9 (ECM deposition) was induced during fibrosis and declined during resolution, while the quiescence program (P1) showed the reciprocal pattern. In the liver, P9 decreased modestly during acute injury but increased sharply in chronic fibrosis, while P1 declined from healthy to acute disease. In skin, most profibrotic programs were induced following injury, whereas P1 showed the opposite pattern. Together, these findings show that the fibroblast programs defined in the cardiac enhancer regulome are conserved across fibrotic diseases and track with disease severity across organs, and the reciprocal relationship between ECM deposition and quiescence emerged as a conserved axis of fibroblast activation in human fibrotic disease.

**Figure 4.**
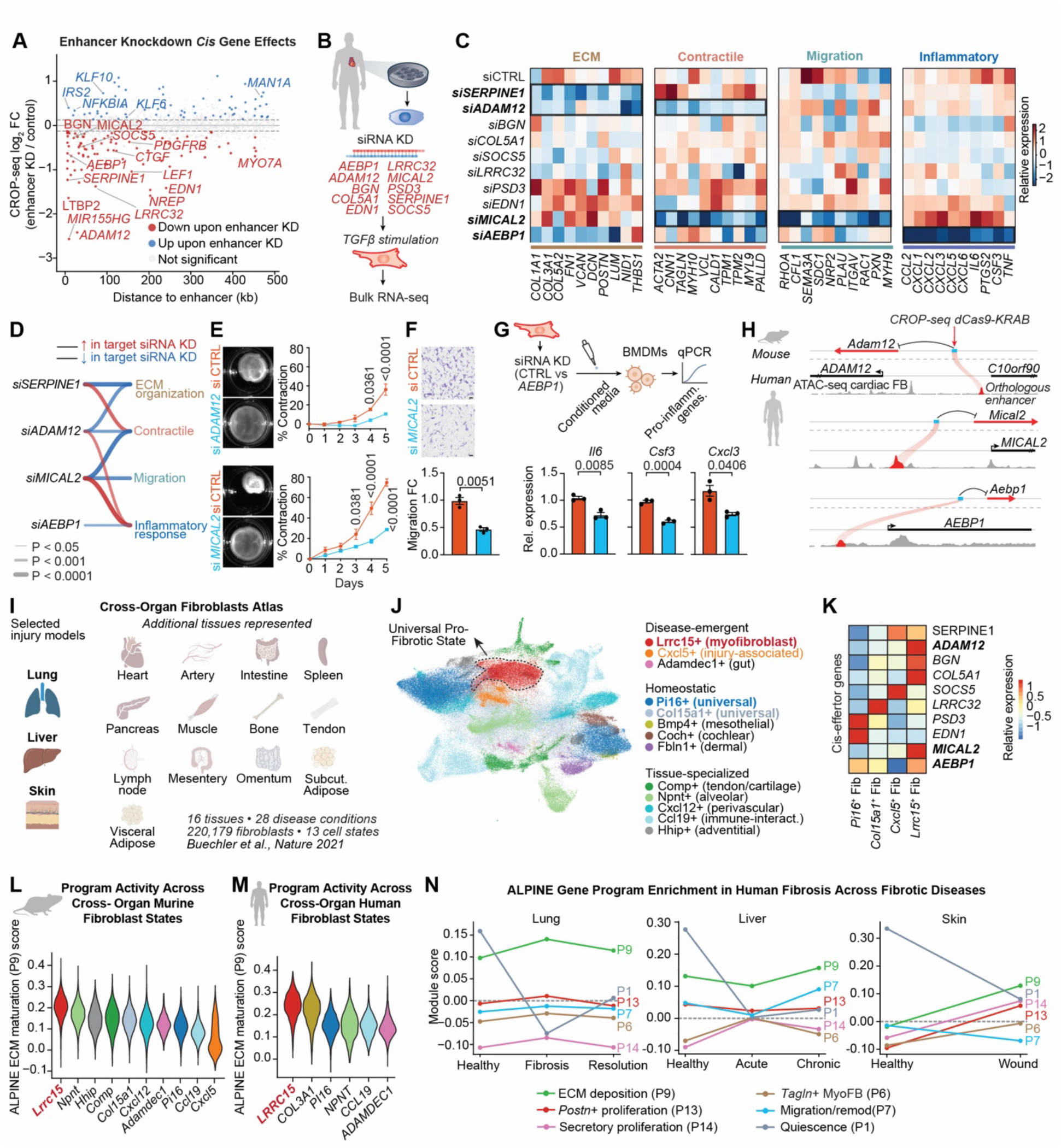
*Cis*-effector genes regulate distinct fibroblast functions and are conserved across organ fibrosis. **A.** CROP-seq cis gene effects (log_2_FC, enhancer KD vs. control) as a function of distance to enhancer. Genes downregulated (red) or upregulated (blue) upon enhancer repression. **B**. Schematic of siRNA knockdown of cis-regulated effector genes (*AEBP1, ADAM12, MICAL2, BGN, COL5A1, SERPINE1, LRRC32, PSD3, EDN1, SOCS5*) in TGFβ-treated human cardiac fibroblasts followed by bulk RNA-seq. **C**. Heatmaps of relative expression changes across siRNA conditions for ECM, contractile, migration and inflammatory gene modules. **D.** Transcriptional shifts across functional gene categories upon siRNA depletion. Lines drawn where P < 0.05 (Mann-Whitney U); thickness proportional to significance. Blue, downregulated; red, upregulated. **E.** Collagen contraction assay images and quantification for *siADAM12* (top) and *siMICAL2* (bottom) versus siCTRL. Representative images at day 5. Mean ± SEM, n=3; two-way ANOVA. **F.** Migration assay images and quantification for *siMICAL2* versus *siCTRL*. Scale bar = 100µm. Mean ± SEM, n=3; unpaired t-test with Welch’s correction. **G.** Schematic of conditioned media experiment (top). Fibroblasts treated with *siCTRL* or *siAEBP1*, conditioned media applied to BMDMs, and RT-qPCR quantification of *Il6*, *Csf3* and *Cxcl3* (bottom). Mean ± SEM, n=3; unpaired t-test with Welch’s correction. **H.** Mouse-to-human conservation at the *Adam12/ADAM12*, *Mical2/MICAL2* and *Aebp1/AEBP1* loci showing CROP-seq dCas9-KRAB perturbation and human cardiac fibroblast ATAC-seq with orthologous enhancers highlighted. **I.** Cross-organ fibroblast atlas spanning 16 tissues, 28 disease conditions and 220,179 fibroblasts across 13 cell states^31^ **J.** UMAP of cross-organ fibroblast states. A universal profibrotic state (*Lrrc15*+ myofibroblast) is highlighted among disease-emergent, homeostatic and tissue-specialized populations. **K.** Relative expression of *cis*-effector genes across fibroblast states (*Pi16*+, *Col15a1*+, *Cxcl5*+ and *Lrrc15*+ fibroblasts). **L,M.** ALPINE ECM maturation (P9) program activity across murine (L) and human (M) cross-organ fibroblast states. **N.** Selected ALPINE gene program enrichment across human fibrotic diseases in lung, liver and skin.

### A compact regulatory module governs a disease-associated cellular state

Cellular responses to injury are governed by complex regulatory networks and are unlikely to depend on a single gene or enhancer. We therefore asked whether a small number of enhancers could account for a large fraction of the profibrotic transcriptional program. We defined a fibrosis gene signature comprising fibroblast genes dynamically regulated during murine heart failure^7^ and also expressed in human cardiac fibroblasts in chronic heart failure or acute myocardial infarction^11^ (n = 110 genes), and used the CROP-seq data to determine how many enhancers must be repressed to suppress increasing fractions of this signature. Repression of just six enhancers, spanning different biological functions including migration, ECM deposition, contractility, proliferation, and mitosis, was predicted to silence 79% of this signature (**Fig. 5A**). We next asked whether simultaneous repression of these six enhancers is sufficient to correct the profibrotic regulatory network and restore fibroblasts to a quiescent state. To enable multiplexed repression, we generated a cardiac fibroblast line expressing dCas12a-KRAB (**Fig. S9A)**, which processes multiple guide spacers from a single chained CRISPR array. We first validated the system by simultaneously targeting the promoters of four genes, *Postn*, *Dap*, *Tfg*, and *Ctgf*, with two guides each, confirming concurrent repression of all four targets (**Fig. S9B**). Having established that the platform supports multiplexed silencing, we delivered an array encoding twelve guide RNAs, two per enhancer, to simultaneously repress the six selected enhancers (dCas12a-6Enh), alongside a matched control carrying ten non-targeting guides (dCas12a-NTG) (**Fig. 5B**). To assess the specificity of enhancer silencing, we profiled the genome-wide chromatin landscape of the dCas12a-6Enh line. H3K9me3 CUT&RUN confirmed strong and selective deposition of repressive chromatin at the six targeted enhancers (**Fig. 5C, Fig. S9C**). Reciprocally, genome-wide ATAC-seq showed that the six enhancers were among the most strongly closed regulatory elements, demonstrating remarkable specificity of the multiplexed system (**Fig. 5D**). Three of the six enhancers were linked by E2G mapping to clear *cis*-regulated target genes: E1 to *Nrep*, a small cytoplasmic protein that modulates cytoskeletal signaling; E4 to *Tgfbi*, a secreted ECM glycoprotein involved in cell adhesion; and E6 to *Rbpj*, a TF with broad roles in cell fate and differentiation. Neither *Nrep* nor *Rbpj* had a previously defined role in cardiac fibroblast activation, but all three were among the genes repressed by *Meox1* deletion *in vivo* (**Fig. 3P**), indicating that the six-enhancer module intersects directly with the MEOX1-regulated profibrotic network. At each of these loci, dCas12a-mediated H3K9me3 deposition at the targeted enhancer was accompanied by reduced chromatin accessibility and decreased expression of the linked gene (**Fig. 5E**). The remaining three targeted enhancers (E2, E3, and E5) showed chromatin silencing at the targeted loci (**Fig. S9D**), but lacked clear *cis*-regulated target genes, suggesting that they may act through more distributed cis-regulatory mechanisms or through trans effects.

To determine how six-enhancer repression rewires the profibrotic regulatory network, we first scored the expression changes across the gene programs defined by the cardiac enhancer regulome. The ECM deposition program (P9) was the most strongly suppressed, followed by the *Myl9*-positive contractile myofibroblast (P10) and *Tagln*-positive contractile myofibroblast (P6) programs (**Fig. 5F, Fig. S9E)**, consistent with a broader loss of chromatin accessibility across the profibrotic enhancer landscape (**Fig. S9F**). Conversely, the quiescence program (P1) was significantly induced (**Fig. 5F**). Multiplexed enhancer repression therefore does not merely dampen individual targets but corrects the profibrotic regulatory network, redirecting fibroblasts toward a quiescent transcriptional state. To understand the breadth of this regulatory rewiring, we integrated genome-wide chromatin accessibility and gene expression changes. Genes linked to enhancers with reduced accessibility were predominantly downregulated, with 55% supported by direct enhancer-to-gene links, compared with only 13% of genes linked to regions with increased accessibility (**Fig. 5G**). The repressed gene set extended beyond the six directly targeted loci to include *Serpine1*, *Tnc*, *Twist2*, *Edn1*, and multiple other regulators of ECM remodeling, migration, and paracrine signaling (**Fig. 5G–H**). Gene regulatory network analysis showed that repression of six enhancers was sufficient to collapse the broader profibrotic transcriptional program (**Fig. 5I**). Critically, many of the downregulated genes, including a broad set of fibroblast-activation TFs, had linked enhancers that lost accessibility in the dCas12a-6Enh line, indicating that the repressive signal propagates through the *cis*-regulatory landscape. Among these, known TF-target relationships trace the cascade throughout the network: *Tead4*, *Twist2*, *Foxd1*, and *Foxf2* converge on the matricellular gene *Ltbp2*; *Creb5* and *Nfatc2* regulate the paracrine factor *Pdgfb*; *Fosl1* and *Etv1* target the migration regulator *Dlc1*; *Atf3* and *Maff* feed into the ERK-AP-1 feedback loop through *Phlda1*; and *Mybl2* and *E2f2* drive the cell-cycle gene *Plk4*. Beyond these defined interactions, ECM, paracrine, migration, and proliferation genes were broadly suppressed, often accompanied by reduced accessibility at their linked enhancers. Together, these changes reveal a coordinated collapse of the profibrotic network, suggesting that a small set of enhancers can sustain the activated fibroblast state through a self-reinforcing regulatory architecture.

Having established that multiplexed enhancer repression collapses the profibrotic regulatory network, we asked whether this translates into functional consequences for fibroblast behavior in disease-relevant contexts. In trans-well migration assays, TGF-β stimulation strongly induced migration in control dCas12a-NTG fibroblasts, but this response was completely abolished in the dCas12a-6Enh line (**Fig. 5J**). Collagen-gel contraction was also impaired, with multiplexed enhancer repression showing a marked reduction in contractile capacity (**Fig. 5K**), accompanied by reduced α-smooth muscle actin expression (**Fig. S9G**). Fibroblasts are known to influence neighboring cells within the tissue microenvironment, and the suppression of paracrine signaling genes upon multiplexed enhancer repression (**Fig. 5I**) suggested that this intervention may alter intercellular communication. To test this, we transferred conditioned medium from TGF-β-stimulated six-enhancer-repressed fibroblasts to bone marrow-derived macrophages and profiled the macrophage transcriptome by bulk RNA-seq. Macrophages exposed to conditioned medium from enhancer-repressed fibroblasts showed broad suppression of cytokine signaling, inflammatory response, and NF-κB pathways, including *Tnf*, *Il1b*, *Ccl2*, *Ccl3*, *Cxcl1*, and *Nos2* (**Fig. 5L, Fig. S9H,I**). Multiplexed enhancer repression therefore not only corrects intrinsic fibroblast pathology but also alters the fibroblast secretome, attenuating macrophage inflammatory activation.

**Figure 5.**
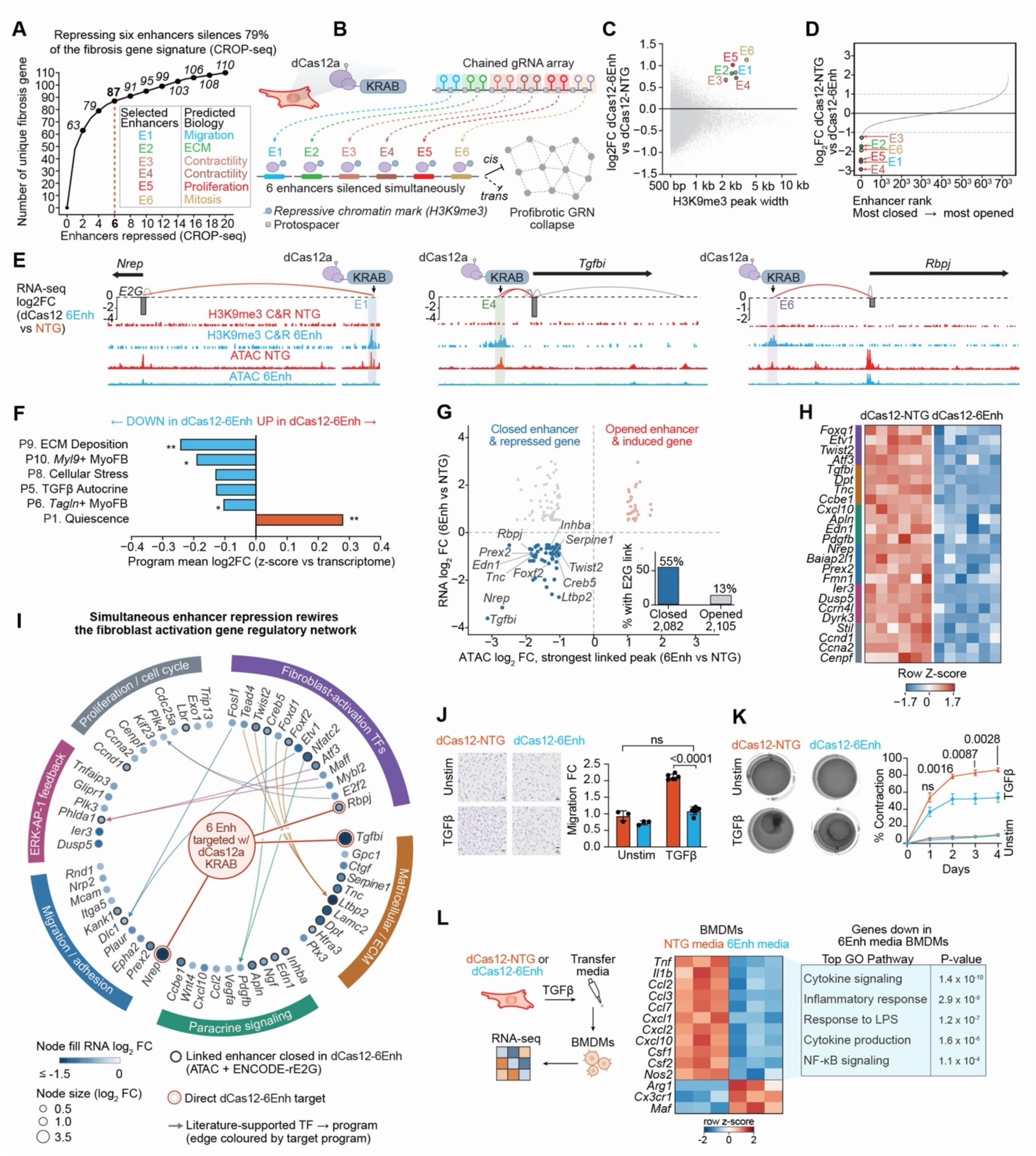
A minimal set of enhancers sustains the profibrotic gene regulatory network in cardiac fibroblasts. **A.** Maximum cumulative number of unique fibrosis signature genes silenced with increasing enhancer repression in CROP-seq. Six enhancers (E1–E6) selected for simultaneous targeting with predicted biological functions annotated. **B.** Schematic of multiplexed dCas12a-KRAB strategy. Chained gRNA array targets six enhancers simultaneously. **C.** Genome-wide H3K9me3 deposition (log_2_FC, dCas12-6Enh vs. dCas12-NTG) as a function of H3K9me3 peak width. The six targeted enhancers (E1–E6) are highlighted. **D.** Ranked enhancer accessibility changes (log2 FC, dCas12-NTG vs. dCas12-6Enh) across all enhancers; E1–E6 highlighted. **E.** Genome browser tracks at the *Nrep*, *Tgfbi* and *Rbpj* loci showing RNA-seq log2 FC (dCas12-6Enh vs. NTG), H3K9me3 CUT&RUN and ATAC-seq in NTG and 6Enh conditions. E2G links indicated. **F.** ALPINE program mean log2 FC (z-score vs. transcriptome) in dCas12-6Enh versus dCas12-NTG. Selected programs shown. **G.** ATAC-seq log₂FC versus RNA-seq log₂FC (6Enh vs. NTG) at enhancer-linked loci. Bar chart (inset): percentage of genes with a direct E2G link among those with closed (55%) versus opened (13%) enhancers. **H.** Heatmap of selected differentially expressed genes between dCas12-NTG and dCas12-6Enh. **I.** Gene regulatory network reconstruction. Simultaneous enhancer repression rewires the fibroblast activation network. Node size and fill reflect RNA log2 FC; edges colored by target program. Black-outlined nodes, linked enhancers closed in dCas12-6Enh (ATAC + ENCODE-rE2G); red-outlined nodes, direct dCas12-6Enh targets; arrows, literature-supported TF-to-target gene connections (colored by target program). **J.** Migration assay images and quantification in dCas12-NTG and dCas12-6Enh fibroblasts in unstimulated and TGFβ conditions. Scale bar = 100µm. Mean ± SEM, n=4; one-way ANOVA with Šídák’s multiple comparisons test. **K.** Collagen contraction assay images and quantification in dCas12-NTG and dCas12-6Enh fibroblasts in unstimulated and TGFβ conditions. Mean ± SEM, n=6; two-way repeated measures ANOVA with Tukey’s multiple comparisons test. **L.** Conditioned media from dCas12-NTG or dCas12-6Enh TGFβ-treated fibroblasts transferred to BMDMs followed by RNA-seq. Heatmap shows selected downregulated genes in BMDMs exposed to 6Enh-conditioned medium, with top enriched GO pathways.

The regulatory rewiring driven by multiplexed enhancer repression encompasses both direct *cis* mechanisms at the targeted loci and broader trans propagation through the downstream network. To determine how individual *cis*-regulated effector genes contribute to the functional phenotype produced by network collapse, we depleted *NREP*, *TGFBI*, and *RBPJ* individually by siRNA in primary human cardiac fibroblasts and profiled transcriptional and functional consequences (**Fig. S10A-C**). Bulk RNA-seq revealed that each depletion affected a distinct transcriptional program: *NREP* depletion strongly suppressed cell cycle and proliferation genes, *TGFBI* depletion preferentially reduced contractile and migratory gene expression, and *RBPJ* depletion suppressed inflammatory signaling genes (**Fig. S10D-F**). Functionally, all three depletions impaired collagen-gel contraction and migration, with *TGFBI* depletion producing the largest reduction in contractile capacity (**Fig. S10G,H**). Thus, these three *cis*-regulated effector genes account for distinct components of the functional phenotype produced by network collapse.

These results establish that a complex, multifunctional disease-associated cellular state can be sustained by a remarkably compact regulatory module, and that its targeted disruption is sufficient to collapse the pathological network and restore cellular homeostasis.

## DISCUSSION

Tissue fibrosis remains one of the most common consequences of chronic disease^3^. At its core is a maladaptive cell state transition in which fibroblasts acquire a complex pathological identity, yet no current therapy targets this process at the regulatory level. Because this transition is multifunctional and involves the coordinated regulation of hundreds of genes^33^, defining the regulatory logic that sustains it demands systematic perturbation rather than observation alone. Here, by combining CRISPRi-based repression of 280 conserved disease-activated enhancers with single-cell transcriptomics, we constructed a comprehensive regulome that maps enhancers to the gene programs they control, reveals a TF hierarchy centered on MEOX1, and demonstrates that a remarkably compact set of enhancers is sufficient to sustain, and when disrupted, collapse an entire pathological cell state.

The regulome resolved 15 gene programs that collectively define the functional landscape of activated fibroblasts, from ECM deposition and contractility to migration, proliferation, and inflammatory signaling, and assigned each to the specific enhancers whose repression alters its activity. This modular architecture reveals that the profibrotic state is not a monolithic program but a network in which distinct enhancers govern separable biological functions. To identify which of these enhancers are most critical for maintaining the disease state, we used a classification framework that scored how each enhancer’s repression shifted fibroblasts along a disease-to-health axis, defining a core set of profibrotic enhancers. Among the TFs that regulate these elements, MEOX1 exerted the broadest effect, with the strongest impact on the ECM deposition and contractile programs, while RUNX1 selectively governed the migratory-paracrine program and c-FOS broadly affected multiple programs, particularly metabolic activation. Notably, MEOX1 and RUNX1 both operated through widespread chromatin remodeling at profibrotic enhancers, whereas ETV4 strongly altered target gene expression without detectable changes in enhancer accessibility, revealing a mechanistically distinct mode of transcriptional control within the same network. Recent works have shown that RUNX1 regulates cardiac fibrosis^34,35^, and that RUNX1 contributes to macrophage-mediated cardiac recovery^12,36^. Our regulome now places RUNX1 within a defined hierarchy and reveals the specificity of its chromatin effects for the migratory-paracrine program. ETV4, primarily studied as an oncogene in multiple cancers ^37,38^, had no prior role in cardiac fibrosis, and its identification here suggests that the profibrotic network recruits regulators beyond the canonical fibrosis TF repertoire.

MEOX1 emerged as the dominant upstream regulator of the profibrotic enhancer landscape. Our previous work identified the epigenetic reader BRD4 as a critical upstream regulator of *Meox1* activation in cardiac fibroblasts and showed that BET inhibition suppresses *Meox1* expression and fibroblast activation in heart failure^7^. We subsequently found that proinflammatory IL-1β secreted by macrophages in the failing heart activates MEOX1, linking immune-driven inflammation directly to the profibrotic transcriptional program^24^. *MEOX1* expression is absent in the healthy heart and strongly induced in disease-activated fibroblasts in both mouse and human heart failure^7,11,12^, mirroring the dynamics of fibroblast activation itself. Through a new conditional allele, we now demonstrate that fibroblast-specific *Meox1* deletion during pressure-overload heart failure reduces fibrosis, improves cardiac function, and rewires fibroblasts from pathological activation toward quiescence, providing the first *in vivo* evidence from a genetic model that MEOX1 is required for the profibrotic cell state. Critically, the *in vivo* transcriptional changes mapped directly onto the CROP-seq regulome: the MEOX1-regulated profibrotic module in vivo converged on the enhancer-defined ECM deposition program, and the quiescence programs that emerged upon *Meox1* loss corresponded to the homeostatic state identified independently through enhancer perturbation *in vitro*. This illustrates a broader principle: a functionally defined enhancer regulome can serve as a reference framework to interpret the consequences of genetic perturbations *in vivo*, enabling a more precise dissection of which regulatory programs are affected by a given intervention than conventional differential expression analysis alone. This convergence also reinforces MEOX1’s role as a pivot between the profibrotic and quiescent fibroblast states. Consistent with this, direct cardiac reprogramming of myofibroblasts *in v*ivo is accompanied by suppression of *Meox1* expression^39^, suggesting that dismantling the MEOX1-driven regulatory program may be a prerequisite for fibroblasts to exit the pathological state and adopt alternative cell fates.

By mapping the *cis*-regulatory changes induced by each enhancer perturbation, we linked the profibrotic enhancer network to the downstream effector genes that execute the biological functions of the activated state. Depleting ten such effectors individually in primary human cardiac fibroblasts revealed that each controls a distinct component of the activated state, demonstrating that the downstream output of the enhancer network is functionally modular. Each of the three most functionally informative effectors governed a distinct arm of the profibrotic state: *ADAM12* depletion reduced ECM deposition and contractility, defining a cell-autonomous profibrotic function for a gene previously used primarily as a lineage marker of profibrotic fibroblasts^40,41^; *MICAL2* depletion selectively impaired contractility and migration, establishing a role for this actin oxidoreductase^42^ in cardiac fibroblast activation that had not been previously described; and *AEBP1* depletion shut down the inflammatory transcriptional program, extending recent *in vivo* work on *AEBP1* in cardiac fibrosis^43^ to the control of proinflammatory signaling. Thus, a single enhancer regulome fans out through specialized effector genes into separable biological functions, each of which can be independently dismantled.

The enhancers that drive this regulome are themselves conserved: accessible, active regulatory elements at the orthologous loci of *ADAM12*, *MICAL2*, and *AEBP1* were confirmed in primary human cardiac fibroblasts, and many of the *cis*-regulated effector genes were preferentially expressed in the LRRC15-positive universal profibrotic fibroblast population that emerges across tissues and disease contexts^31,44,45^. The gene programs defined by the cardiac regulome extended well beyond the heart: the ECM deposition program showed the highest activity in profibrotic fibroblasts across 16 tissues in both mouse and human, and the reciprocal relationship between ECM deposition and quiescence was dynamically regulated across lung fibrosis, liver fibrosis, and skin wound healing, tracking with disease severity and resolution. Cross-tissue fibroblast atlases have catalogued conserved cell states across organs^31^, but a central unresolved question is how much of the underlying regulatory logic is shared across fibrotic disorders. Our data suggest that the overlap is substantial: the same enhancer-defined programs, controlled by conserved regulatory elements and executed through shared effector genes, govern fibroblast activation across tissues. Tissue-specific microenvironments, cellular niches, and upstream signals will undoubtedly impose distinct regulatory layers, but the conservation of a core enhancer-defined architecture across organs raises the possibility that targeting this shared regulatory logic could complement tissue-specific approaches in fibrotic disease.

The conservation of this regulatory architecture across tissues and diseases raises a further question: can the complexity of the profibrotic network be reduced to a minimal set of regulatory elements whose disruption is sufficient to collapse the pathological state? We took a reductionist approach, reasoning that defining such a module would open the possibility of targeting disease-associated cell states through compact, combinatorial interventions rather than individual genes or pathways. Repression of just six enhancers with a multiplexed dCas12a-KRAB system^46,47^, out of the hundreds active in disease-state fibroblasts, was sufficient to collapse the profibrotic state, suggesting that the regulatory logic maintaining complex pathological cell states may be far more concentrated than their phenotypic complexity implies. Notably, three of the six targeted enhancers govern genes that are also direct MEOX1 targets, suggesting that the minimal module and the MEOX1-driven hierarchy converge on the same core regulatory architecture.

This ability to redirect fibroblasts toward quiescence through targeted regulatory intervention speaks to a broader reconceptualization of fibrosis. Fibrosis was long considered an irreversible endpoint, but lineage tracing in the liver demonstrated that activated myofibroblasts can revert to a quiescent-like phenotype upon cessation of injury^5^, and dynamic fibroblast state transitions have since been mapped in human heart failure recovery^12^. BET inhibition can reverse the profibrotic transcriptional state in cardiac fibroblasts^7^, and the present findings now define the regulatory architecture through which this plasticity operates. This suggests that even in chronic fibrotic disease with established tissue remodeling, the pathological cell state may remain accessible to regulatory intervention, and that strategies targeting the enhancer-level logic rather than individual downstream mediators could offer a path toward reversing, not merely slowing, fibrotic disease.

Together, these findings establish a comprehensive regulatory framework for fibroblast activation in disease, connecting conserved enhancers to gene programs, TFs, effector genes, and ultimately to the cellular behaviors that drive tissue fibrosis. More broadly, this work shows that a complex profibrotic state can be dismantled by targeting a compact regulatory network, providing a conceptual framework for therapeutically reprogramming fibroblast states.

## Acknowledgments

We thank Lin Ye for discussions on lentiviral library preparation, and Ronald J. Vagnozzi, Barbara González-Terán, Alexis Combes, Tomohiro Nishino and Mauro W. Costa for providing feedback. We thank Schuyler Melore (Duke University) for facilitating the material transfer and for valuable technical guidance on the pre-crRNA array cloning. Sequencing for the MPRA was supported by Rebecca Porritt and the SBP Genomics Shared Resource. We thank the Gladstone Flow Cytometry Core and the Gladstone Genomics Core for technical support. We thank Stanford University and the Stanford Research Computing Center for providing computational resources and support as part of the Sherlock High-Performance Compute Cluster.

## Author contribution

M.A. and P.F.P. conceived the project with input from J.M.A. M.A. wrote the manuscript with input from P.F.P. and J.M.A. A.M.Z. performed CROP-seq experiments. Y.L., A. Mahr, and P.F.P. performed CROP-seq computational analysis with input from M.A., J.M.A., C.W., T.Y., B.A., C.-M.L., and S.J. B.C. and S.R. performed MPRA experiments. V.X. and H.Z. generated TF knockdown lines. M.S. and J.M.E. performed E2G analysis. A.M.Z., R.N., H.Z., Y.H., C.Y.L., A. Padmanabhan, Y.A., and K.A. performed Meox1 *in vivo* experiments. M.A. and J.M.A. performed scRNA-seq analysis. A.M.Z., H.Z., V.X., and A. Pelonero performed and analyzed CUT&RUN and ATAC-seq experiments. H.Z. and B.R. performed functional assays in cardiac fibroblasts. V.X. and H.Z. generated the dCas12a system and performed multiplexed repression experiments. M.A., P.F.P., and J.M.A. supervised the study.

---

**Supplementary Figure 1.**
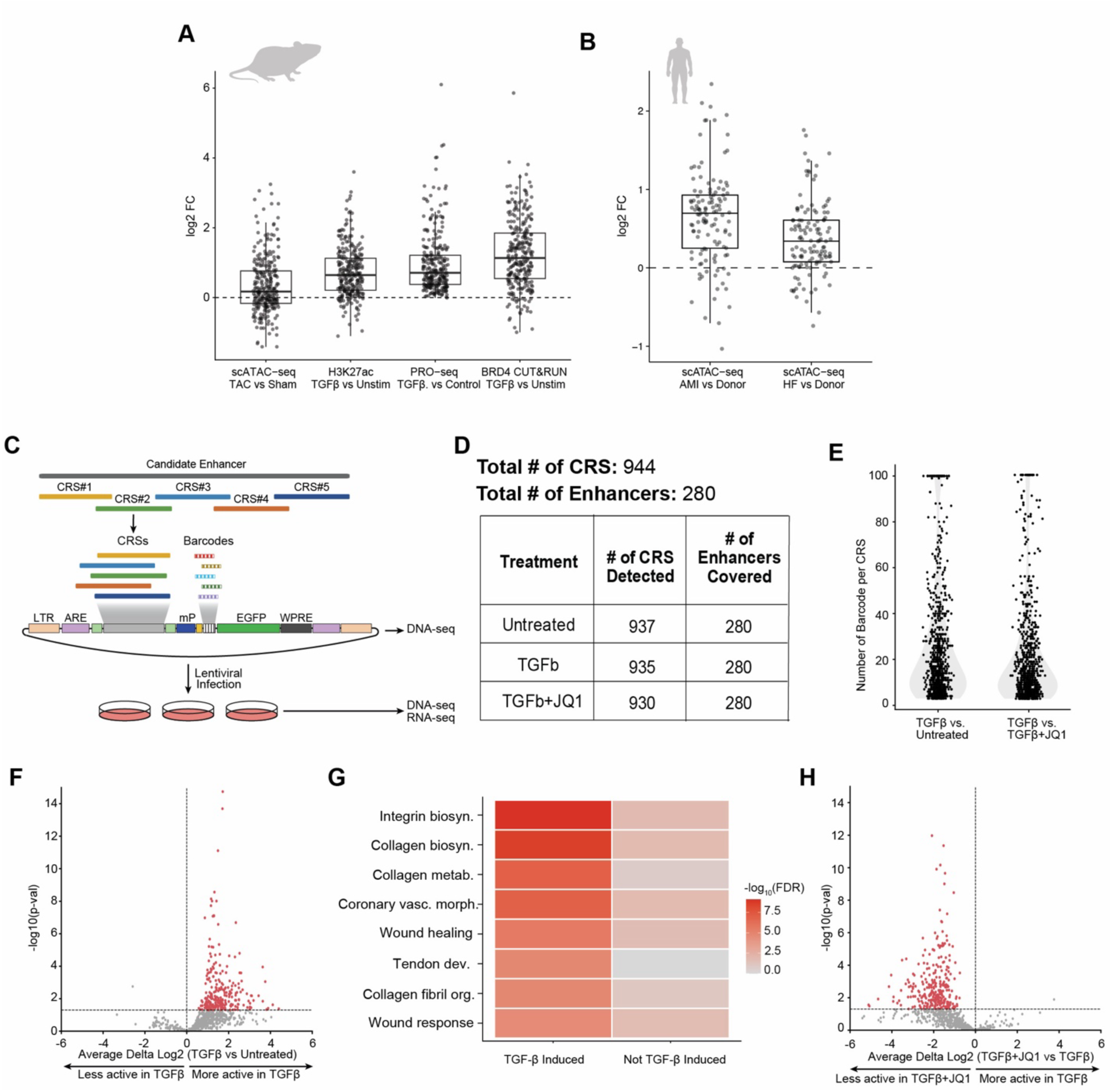
Disease-associated enhancer identification and functional validation by MPRA. **A.** Log_2_ fold change at disease-associated fibroblast enhancers across mouse chromatin assays: *in vivo* scATAC-seq TAC vs. Sham, *in vitro* H3K27ac TGFβ vs. Unstim, *in vitro* PRO-seq TGFβ vs. Unstim, and *in vitro* BRD4 CUT&RUN TGFβ vs. Unstim. **B.** Log_2_ fold change at disease-associated fibroblast enhancers in human cardiac fibroblasts (scATAC-seq AMI vs. Donor and HF vs. Donor). **C.** Schematic of the MPRA workflow. Candidate enhancers are tiled with candidate regulatory sequences (CRSs), each tagged with unique barcodes in a lentiviral reporter construct. **D.** Table showing total number of CRS (944) and enhancers (280) detected across treatment conditions (Untreated, TGFβ, TGFβ+JQ1). **E.** Number of barcodes per CRS in TGFβ vs. Untreated and TGFβ vs. TGFβ+JQ1 comparisons. **F.** Volcano plot of CRS activity (average delta log_2_ TGFβ vs. Untreated). Hits called at adjusted P < 0.05. **G.** GO pathway enrichment for TGFβ-induced enhancers comparing TGFβ-induced and non-TGFβ-induced CRS. **H.** Volcano plot of CRS activity (average delta log_2_ TGFβ+JQ1 vs. TGFβ). Hits called at adjusted P < 0.05.

**Supplementary Figure 2.**
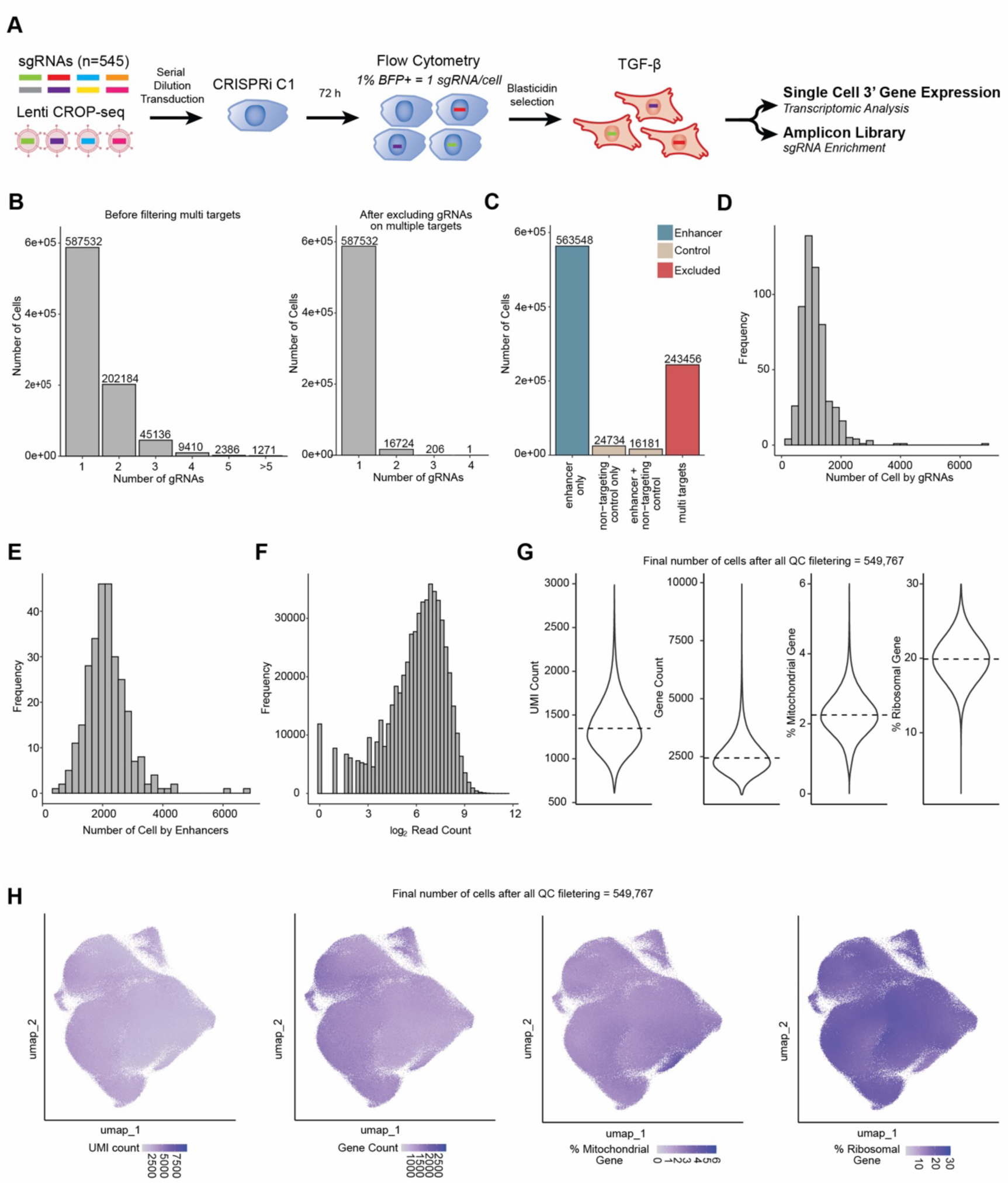
CRISPRi CROP-seq quality control. **A.** Schematic of CROP-seq experimental workflow. sgRNAs (n=545) delivered by serial dilution lentiviral transduction into cardiac fibroblast CRISPRi cells, clone 1 selection, flow cytometry (1% BFP+ = 1 sgRNA/cell), blasticidin selection, TGFβ treatment, followed by single-cell 3’ gene expression for transcriptomic analysis and amplicon library for sgRNA enrichment. **B.** Distribution of the number of sgRNAs detected per cell before (left) and after (right) removal of cells assigned to multiple targets. Cells retained with more than one sgRNA are those in which the multiple sgRNAs target the same enhancer, or are distinct non-targeting control sgRNAs. **C.** Distribution of sgRNA target composition across cells (single-enhancer only, non-targeting control only, single-enhancer + non-targeting control, multi targets). **D.** Histogram of the number of cells assigned per sgRNA. **E.** Histogram of the number of cells assigned per enhancer. **F.** Histogram of sgRNA log_2_ read counts. **G.** Violin plots of UMI count, gene count, and percentage of mitochondrial and ribosomal gene expression per cell in the CROP-seq dataset. After all quality-control filtering, 549,767 cells were retained for downstream analysis. **H.** UMAP plots colored by UMI count, gene count, percentage of mitochondrial genes and percentage of ribosomal genes.

**Supplementary Figure 3.**
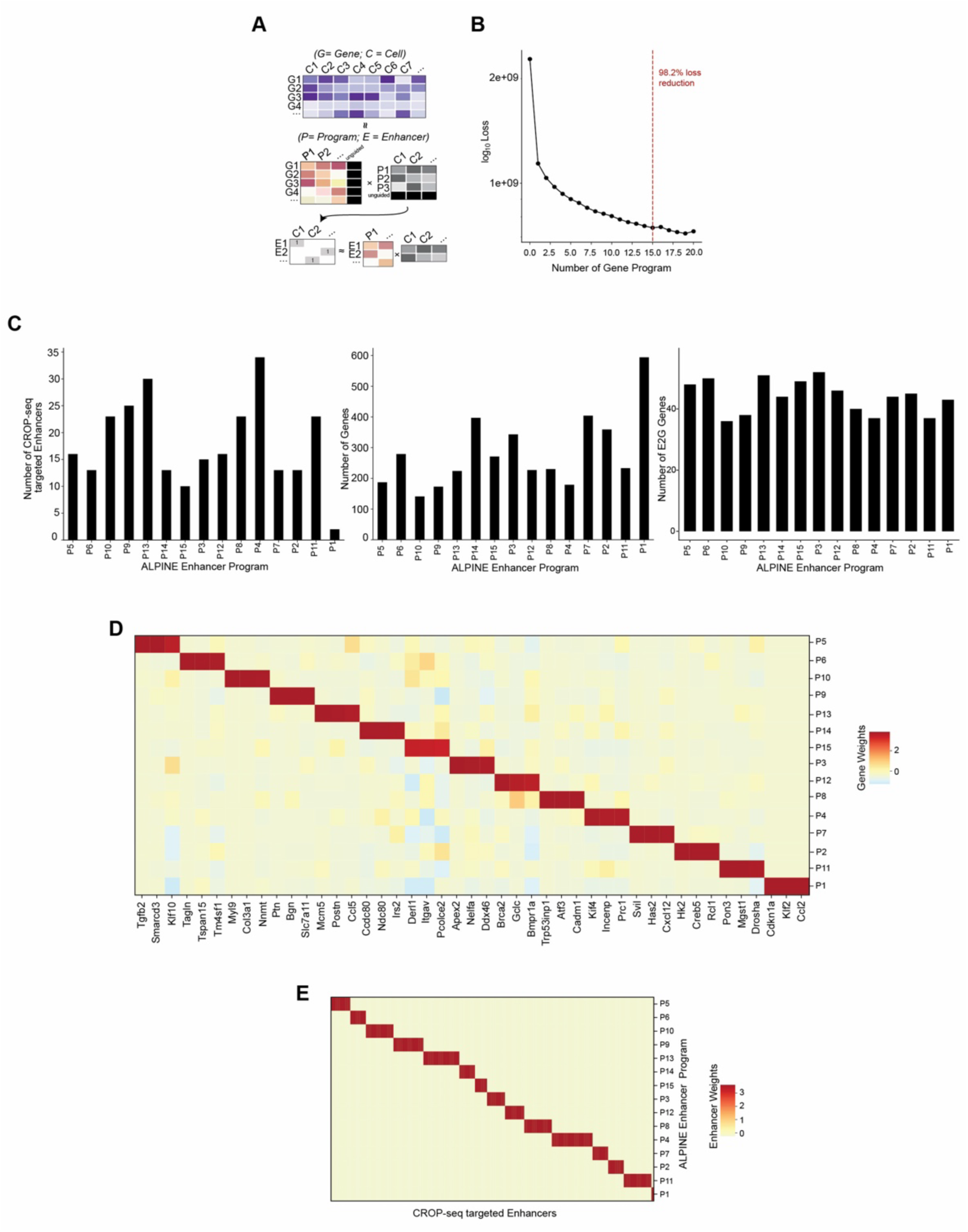
Semi-supervised decomposition of CROP-seq data into fibroblast gene programs. **A.** Schematic overview of the ALPINE framework. Semi-supervised non-negative matrix factorization jointly models gene and cell matrices to define programs linked to enhancers and genes. **B.** Model loss across different numbers of gene programs. The selected number of programs (15) achieved 98.2% of the maximum loss reduction observed across first 20 programs. **C.** Bar plots showing the number of CROP-seq targeted enhancers (left), genes (center) and E2G genes (right) across ALPINE gene programs. **D.** Heatmap of genes displayed in Figure 1F across ALPINE gene programs. **E.** Heatmap of enhancer weights across ALPINE gene programs for all CROP-seq targeted enhancers.

**Supplementary Figure 4.**
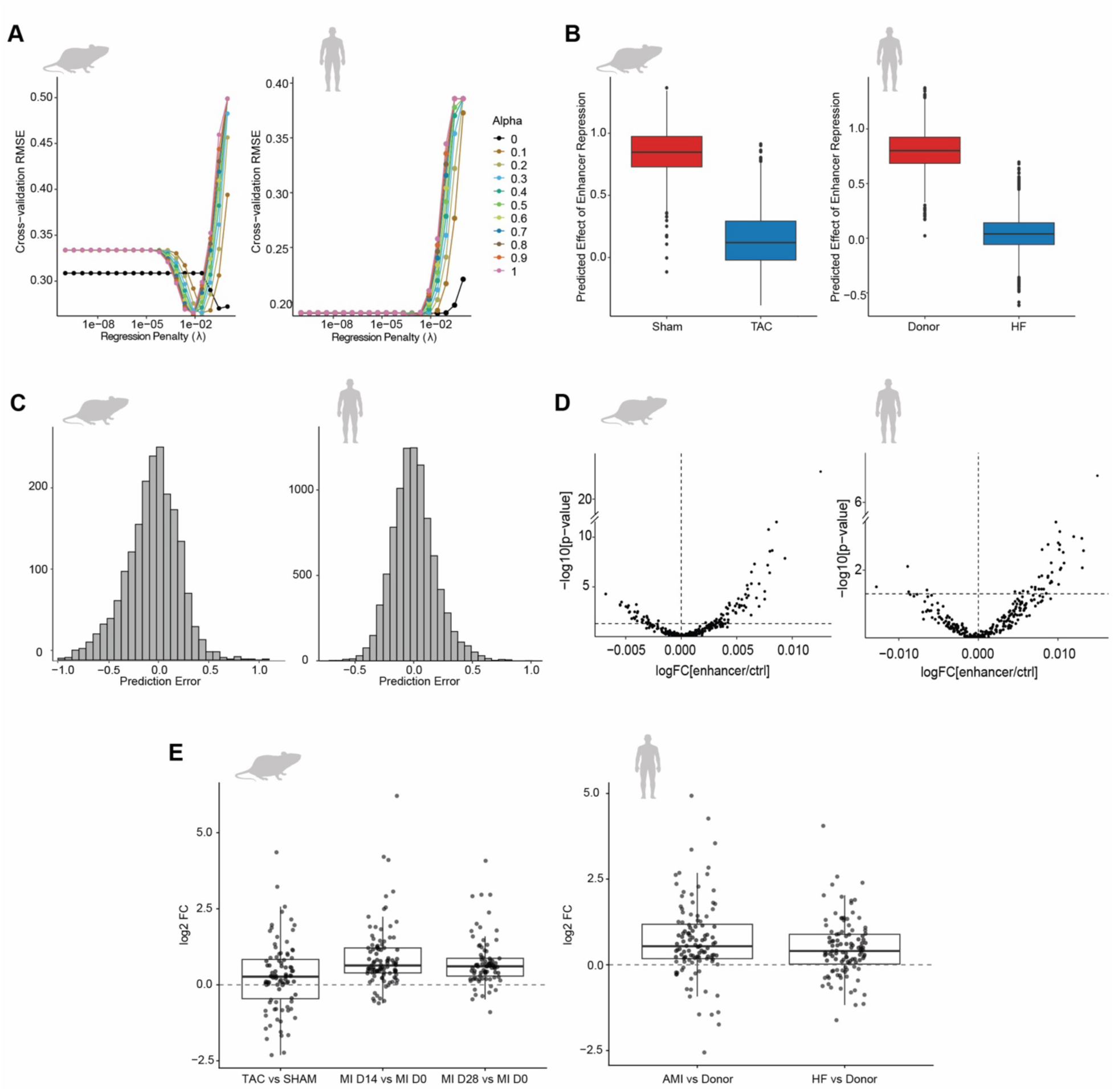
Machine-learning disease classifier identifies profibrotic enhancers across mouse and human models. **A.** Cross-validation RMSE (root mean square error) across regression penalty (λ) for different elastic net mixing parameters (α, colored lines) in the mouse (left) and human (right) logistic regression models. **B.** Predicted effect of enhancer repression in Sham vs. TAC (mouse, left) and Donor vs. HF (human, right). **C.** Distribution of prediction error for the mouse (left) and human (right) models. **D.** Volcano plots of differential enhancer perturbation effects (log_2_ FC enhancer vs. control) in the mouse (left) and human (right) models. **E.** Log_2_ fold change of differentially expressed E2G genes associated with profibrotic enhancers in mouse disease models (TAC vs. Sham, MI D14 vs. D0, MI D28 vs. D0; left) and human (AMI vs. Donor, HF vs. Donor; right).

**Supplementary Figure 5.**
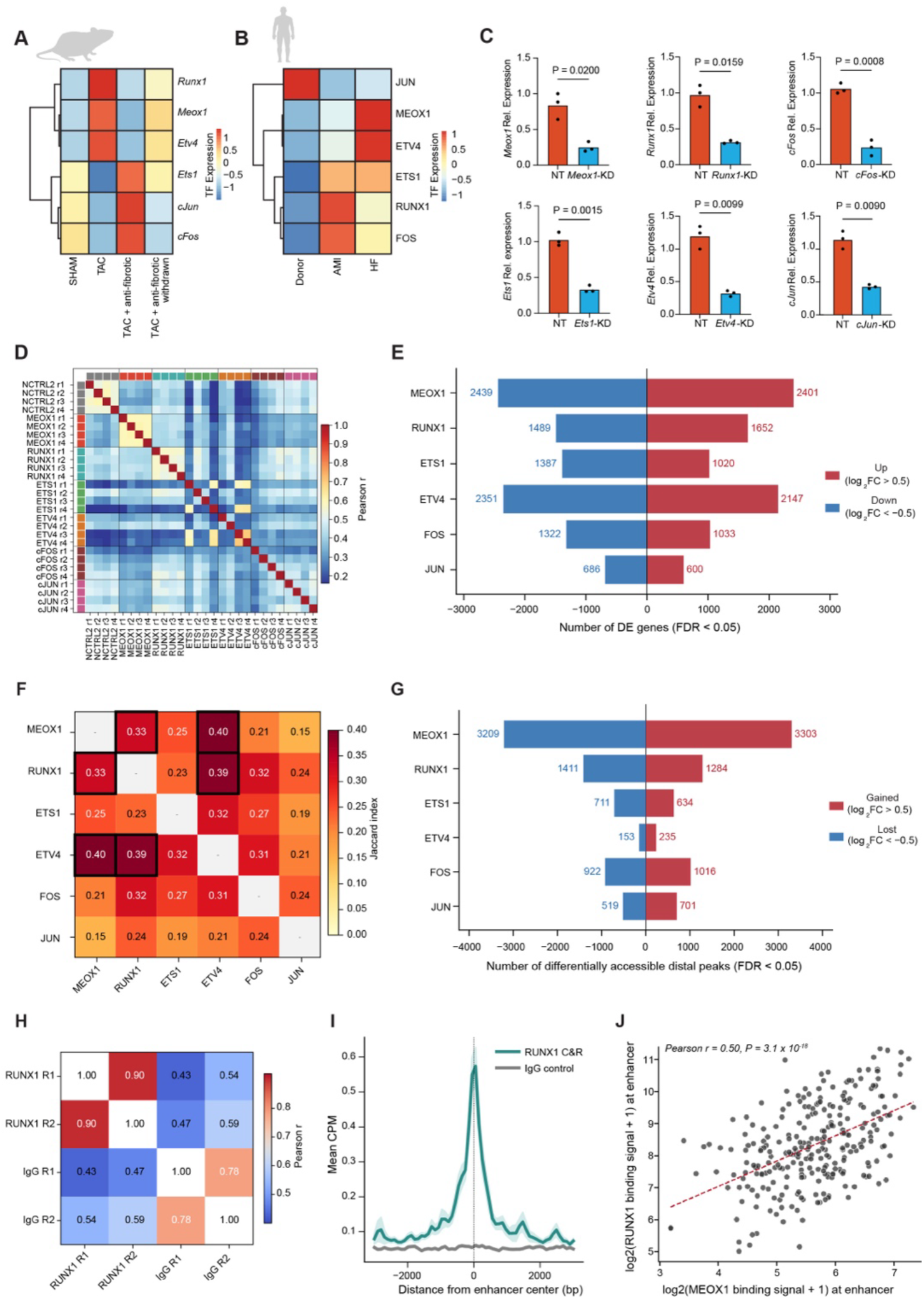
Profibrotic transcription factors regulate distinct gene and chromatin programs. **A.** Heatmap of TF expression in the mouse model of heart failure (Sham, TAC, TAC + anti-fibrotic, TAC + anti-fibrotic withdrawn). **B.** Heatmap of TF expression in human cardiac fibroblasts (Donor, AMI, HF). **C.** RT-qPCR validation of CRISPRi knockdown for each TF (*Meox1*, *Runx1*, *cFos*, *Ets1*, *Etv4*, *cJun*). Mean ± SD, n=3; unpaired t-test. **D.** Sample-to-sample pairwise Pearson correlation between non-targeting control and each TF knockdown RNA-seq replicates. **E.** Number of differentially expressed genes (FDR < 0.05) upon knockdown of each TF. Red, upregulated (log_2_FC > 0.5); blue, downregulated (log_2_FC < −0.5). **F.** Heatmap of pairwise Jaccard index comparing differentially expressed gene sets (FDR < 0.05, |log_2_FC| > 0.5) between TF knockdowns. **G.** Number of differentially accessible distal ATAC-seq peaks (FDR < 0.05) upon knockdown of each TF. Red, gained (log_2_FC > 0.5); blue, lost (log_2_FC < −0.5). **H.** Sample-to-sample pairwise Pearson correlation between RUNX1 CUT&RUN and IgG control replicates. **I.** Metaplot of mean CPM signal for RUNX1 CUT&RUN versus IgG control centered on enhancers. **J.** Correlation between MEOX1 and RUNX1 binding signal at the CROP-seq targeted enhancers (Pearson r = 0.50, P = 3.1 × 10⁻¹⁸).

**Supplementary Figure 6.**
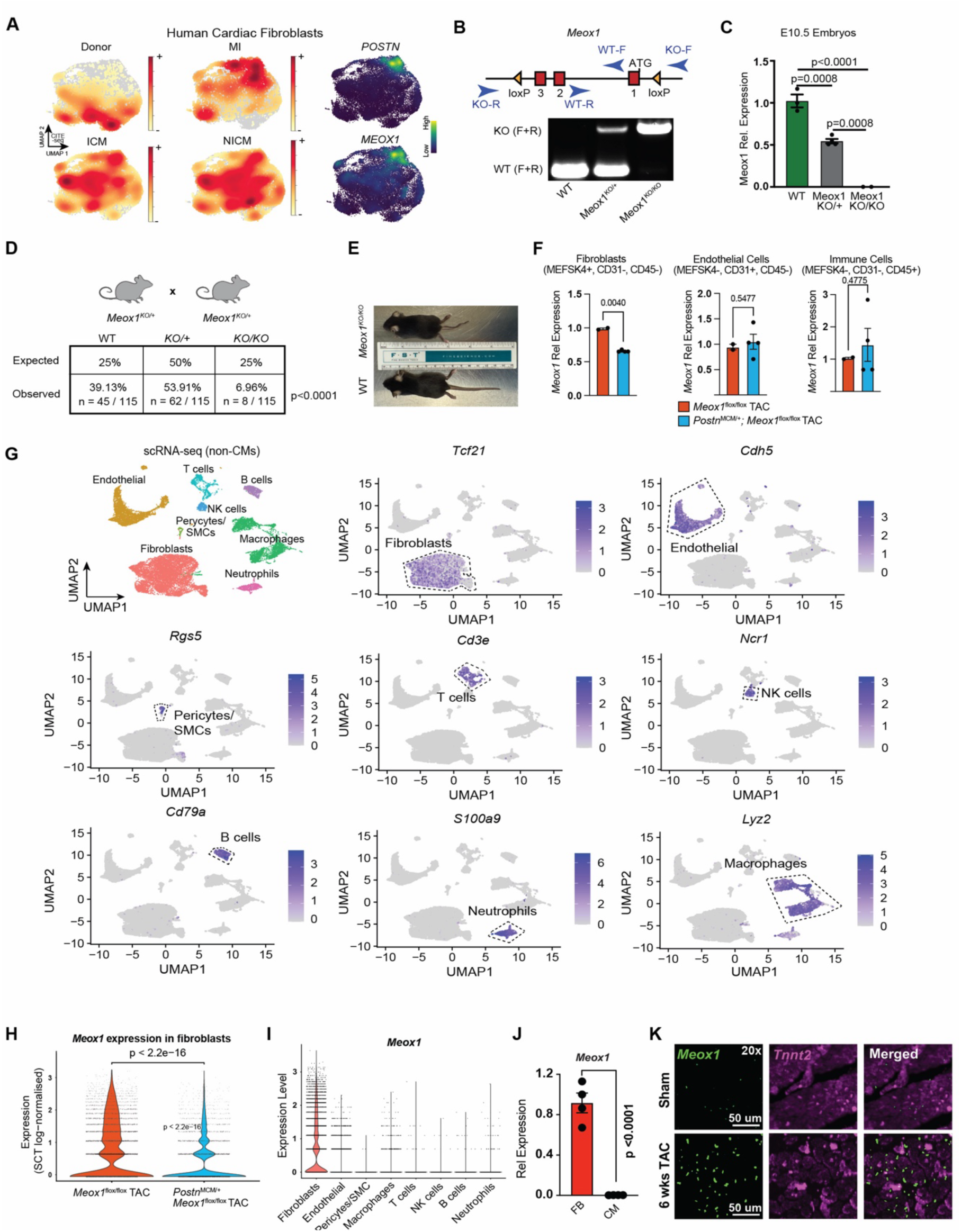
Generation and validation of activated fibroblast-specific *Meox1* conditional knockout mice. **A.** UMAP of human cardiac fibroblasts showing MEOX1 expression overlaps with POSTN-positive fibroblasts and is enriched in heart failure (ICM and NICM). **B.** Recombination strategy of *Meox1*^flox^ animals with positions of genotyping primers (Top). Agarose gel electrophoresis showing WT, *Meox1*^KO/+^ and *Meox1*^KO/KO^ genotypes (bottom). **C.** RT-qPCR validates incremental loss of *Meox1* expression in E10.5 embryos from *Meox1*^KO/+^ × *Meox1*^KO/+^ breeding. One-way ANOVA. **D.** *Meox1*^KO/KO^ animals are not born at expected Mendelian ratios (6.96% observed vs. 25% expected at n=115). Chi-square test, P < 0.0001. **E.** Representative image of *Meox1*^KO/KO^ alongside its WT littermate. **F.** RT-qPCR validation of *Meox1* knockdown in FACS-sorted cell populations (fibroblasts: MESK4+, CD31−, CD45-; endothelial: MESK4−, CD31+, CD45−; immune: MESK4−, CD31−, CD45+) from *Meox1*^flox/flox^ TAC and *Postn*^MCM+^;*Meox1*^flox/flox^ TAC hearts. **G.** Expression feature plots of cell population markers in scRNA-seq of non-cardiomyocyte cells (*Tcf21*, fibroblasts; *Cdh5*, endothelial; *Rgs5*, pericytes/SMCs; *Cd3e*, T cells; *Ncr1*, NK cells; *Cd79a*, B cells; *S100a9*, neutrophils; *Lyz2*, macrophages). **H.** Violin plot of *Meox1* expression in fibroblasts from *Meox1*^flox/flox^ TAC and *Postn*^MCM+^;*Meox1*^flox/flox^ TAC hearts. Wilcoxon rank-sum test, P < 2.2e−16. **I.** Violin plot of *Meox1* expression across cell types 6 weeks post-TAC. **J.** *Meox1* is expressed specifically in fibroblasts and not cardiomyocytes as quantified by RT-qPCR. Unpaired t-test with Welch’s correction, P < 0.0001. **K.** RNAscope of *Meox1* (green) and *Tnnt2* (magenta) in Sham and 6-week TAC hearts. *Meox1* is upregulated after TAC and does not overlap with cardiomyocytes (*Tnnt2*). Scale bar = 50 µm.

**Supplementary Figure 7.**
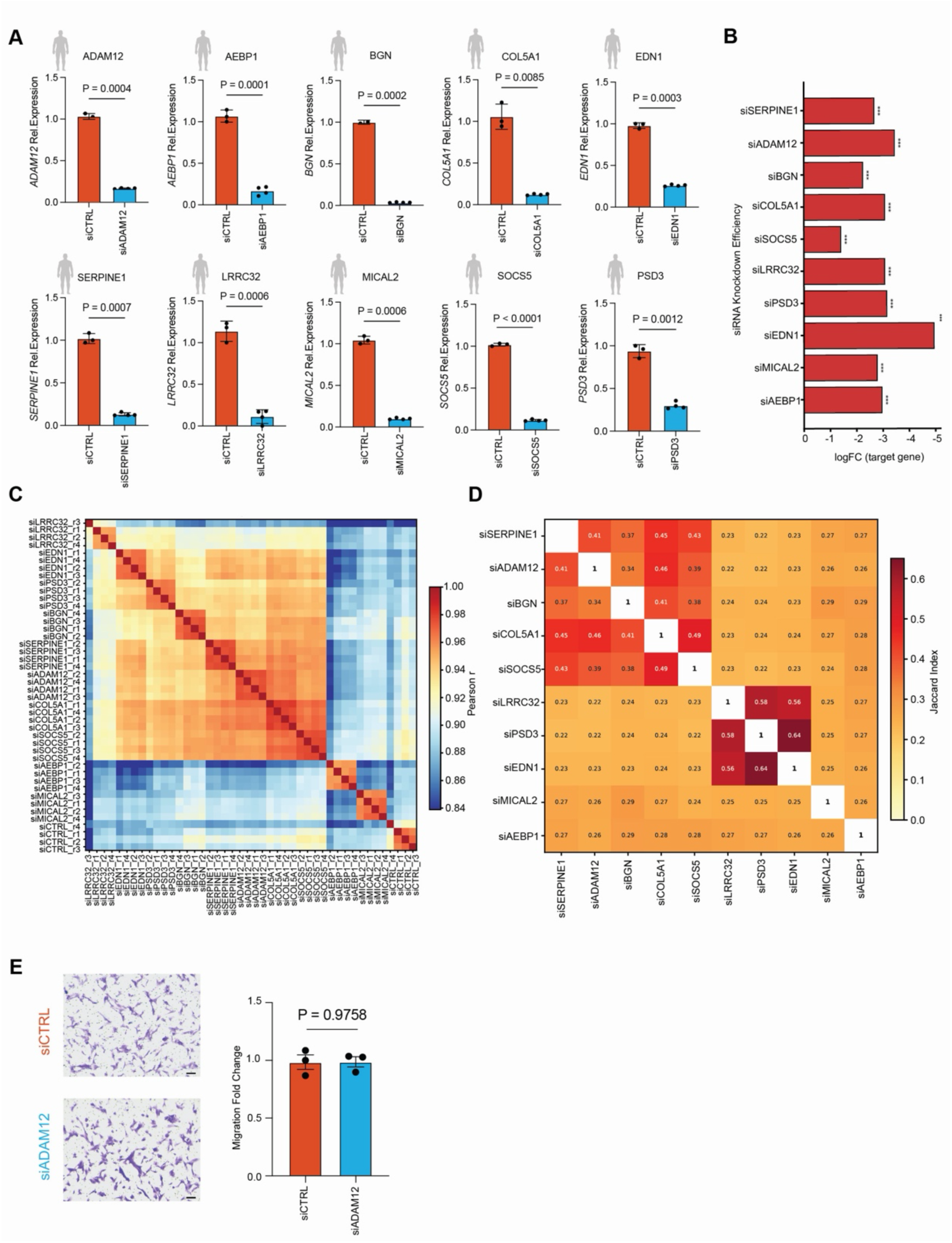
Cis-regulated effector gene knockdown validation in human cardiac fibroblasts. **A.** RT-qPCR validation of siRNA knockdown for *ADAM12*, *AEBP1*, *BGN*, *COL5A1*, *EDN1*, *SERPINE1*, *LRRC32*, *MICAL2*, *SOCS5* and *PSD3* in primary human cardiac fibroblasts. Mean ± SD; n = 3 (siCTRL), n = 4 (each siRNA target); unpaired t-test. **B.** siRNA knockdown efficiency (log_2_FC of target gene by RNA-seq) for each siRNA relative to siCTRL. **C.** Sample-to-sample pairwise Pearson correlation (genome-wide) between individual RNA-seq replicates across all siRNA conditions. **D.** Heatmap of pairwise Jaccard index comparing differentially expressed gene sets (FDR < 0.05, |log_2_FC| > 0.5) between siRNA knockdowns. **E.** Representative transwell migration assay images and quantification for si*ADAM12* relative to siCTRL. Mean ± SEM, n=3; unpaired t-test with Welch’s correction.

**Supplementary Figure 8.**
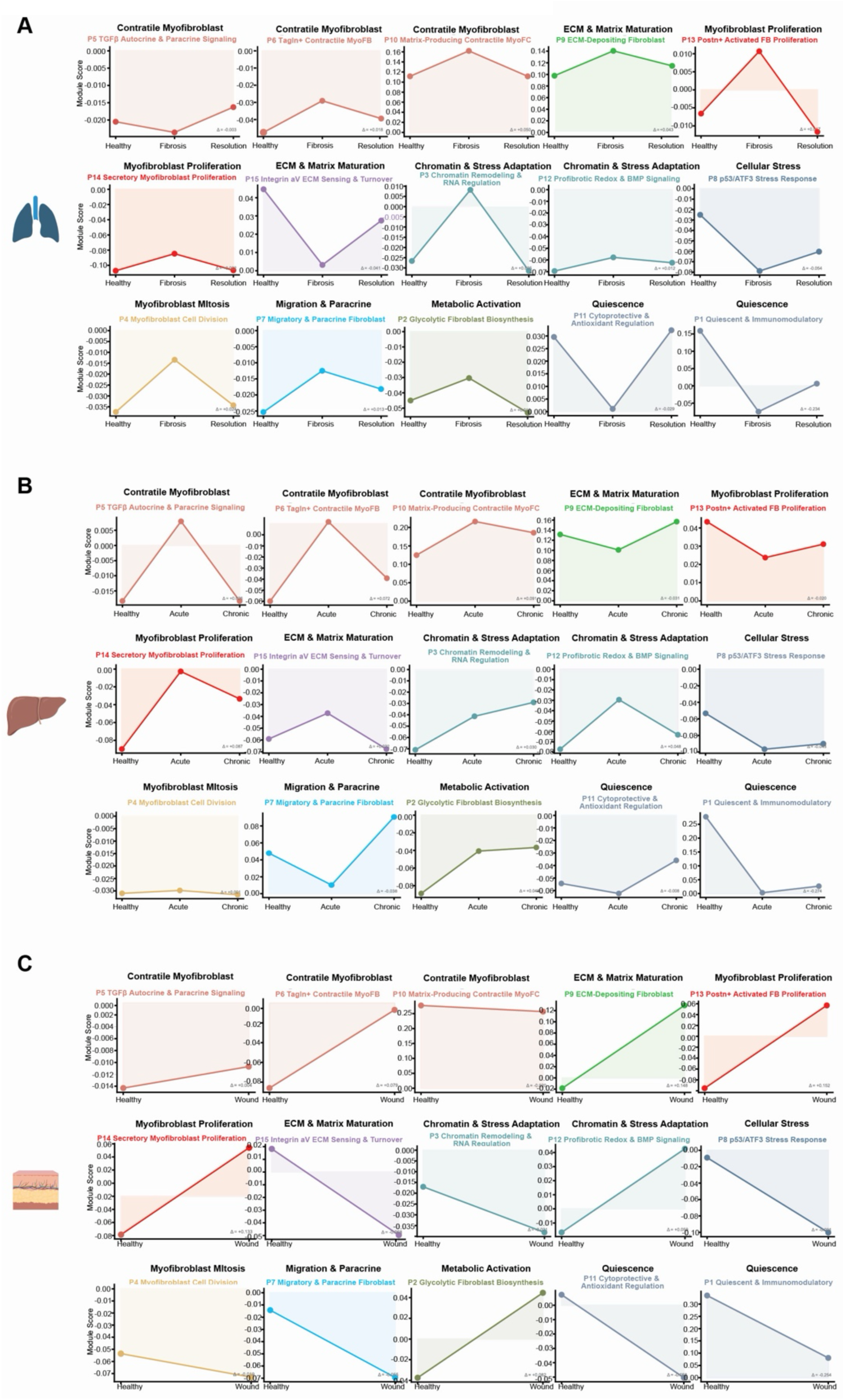
ALPINE gene program conservation across organ fibrosis. **A.** Module scores for ALPINE gene programs across lung fibrosis disease stages (Health, Fibrosis D11, Resolution D21). **B.** Module scores for ALPINE gene programs across liver fibrosis disease stages (Health, Acute, Chronic). **C.** Module scores for ALPINE gene programs across skin fibrosis (Health, Wound).

**Supplementary Figure 9.**
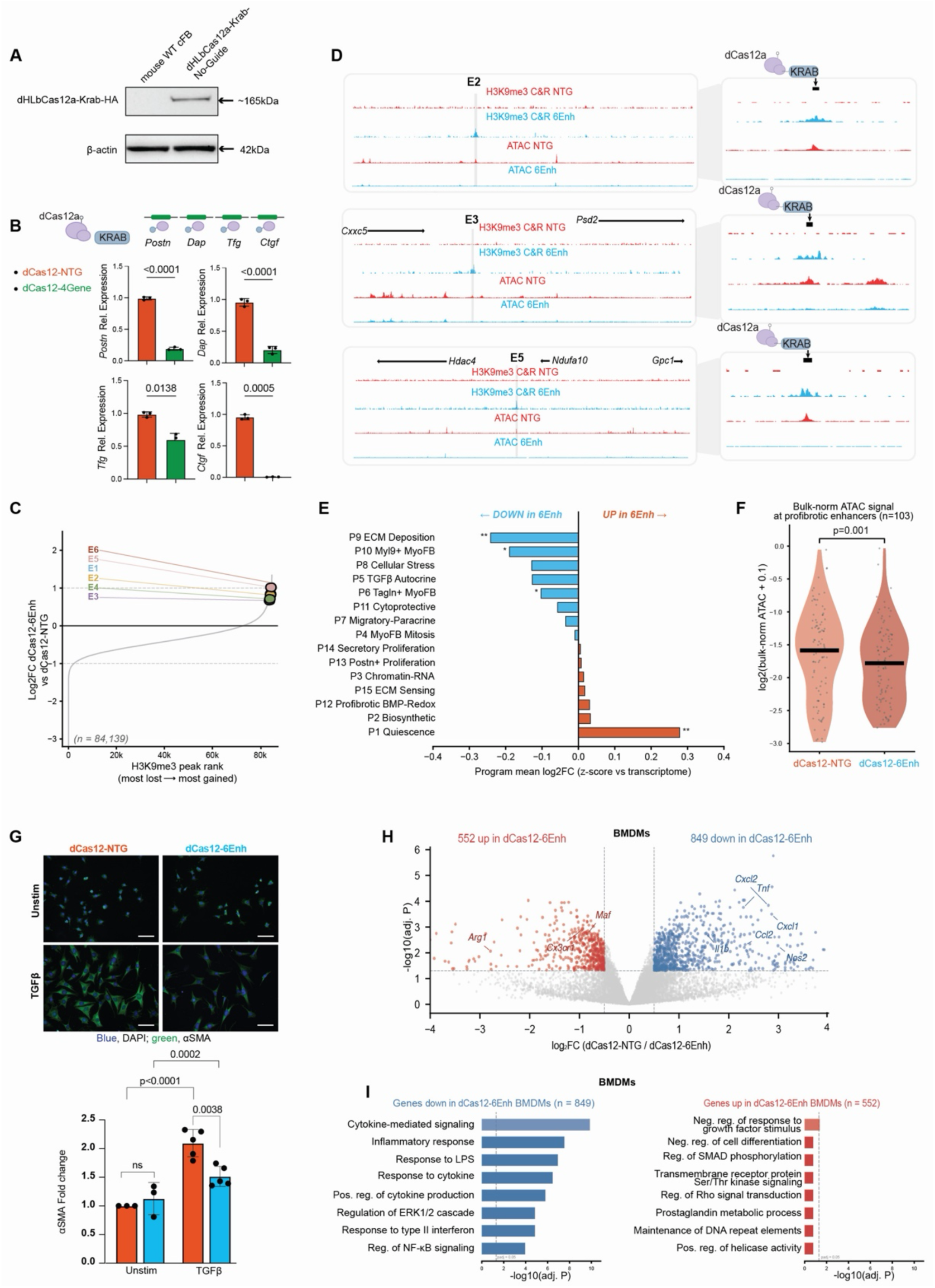
Validation of multiplexed dCas12a-KRAB enhancer silencing in cardiac fibroblasts. **A.** Western blot validation of HA-tagged dHLbCas12a-KRAB protein expression (∼165 kDa) in transduced mouse cardiac fibroblasts. β-actin (∼42 kDa) as loading control. **B.** Schematic of dCas12a targeting of gene TSS (top). RT-qPCR of *Postn*, *Dap*, *Tfg* and *Ctgf* in cardiac fibroblasts expressing dHLbCas12a-KRAB with non-targeting (dCas12-NTG) or 4-gene-targeting guides (dCas12-4Gene). Mean ± SD, n=3; unpaired t-test. **C.** Rank plot of log_2_FC in H3K9me3 signal (dCas12-6Enh vs. dCas12-NTG) across all H3K9me3 peaks (n=84,139). The six targeted enhancers (E1–E6) are highlighted. **D.** Genome browser tracks of H3K9me3 CUT&RUN and ATAC-seq signal at enhancers E2, E3, and E5 in dCas12a-NTG (red) and dCas12a-6Enh (blue) cardiac fibroblasts. Right panels show magnified views of each targeted enhancer. **E.** ALPINE z-score log_2_FC per program between dCas12-NTG and dCas12-6Enh. *FDR < 0.05, **FDR < 0.01. **F.** Bulk-normalized ATAC-seq signal at profibrotic (Q1) enhancers (n=103) in TGFβ-stimulated cardiac fibroblasts expressing dCas12-NTG or dCas12-6Enh. P = 0.001. **G.** Representative immunofluorescence images of αSMA (green) and nuclei (blue) in cardiac fibroblasts expressing dCas12-NTG or dCas12-6Enh, unstimulated or treated with TGFβ. Scale bar = 100 µm. Quantification of αSMA fold change; mean ± SD, n=3 (Unstim) or n=5 (TGFβ); one-way ANOVA. **H.** Volcano plot of differentially expressed genes in BMDMs treated with conditioned media from dCas12-NTG vs. dCas12-6Enh fibroblasts. 552 genes upregulated, 849 genes downregulated (adj. P < 0.05, |log2FC| > 0.5). **I.** GO enrichment of genes downregulated (n=849, left) and upregulated (n=552, right) in dCas12-6Enh BMDMs, ranked by −log10(adj. P).

**Supplementary Figure 10.**
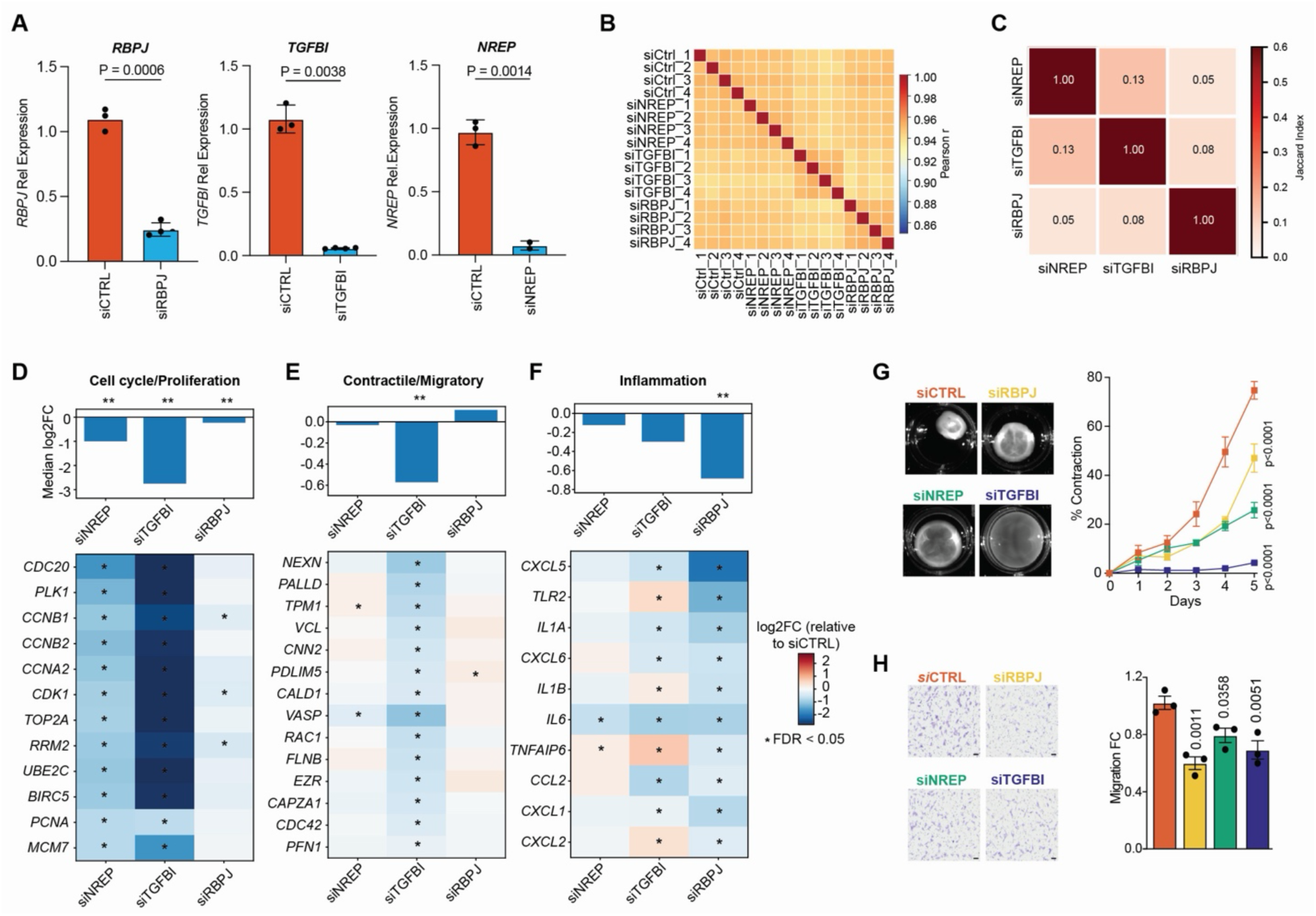
Transcriptional and functional consequences of Cas12 *cis*-regulatory target gene knockdown in primary human cardiac fibroblasts. A. RT-qPCR validation of siRNA knockdown of *RBPJ*, *TGFBI* and *NREP* in primary human cardiac fibroblasts. Mean ± SD, n=3; unpaired t-test. B. Sample-to-sample Pearson correlation of voom-normalized log_2_CPM values (n=4 per condition). All pairwise correlations r > 0.86. C. Jaccard index of differentially expressed gene sets between siRNA conditions (FDR < 0.05, |log_2_FC| > 0.5). D. Median log_2_FC and gene-level heatmap for the cell cycle/proliferation program across si*NREP*, si*TGFBI* and si*RBPJ* conditions. Module-level significance by Wilcoxon signed-rank test with Benjamini–Hochberg correction. Asterisks within heatmaps denote FDR < 0.05. E. Median log_2_FC and gene-level heatmap for the contractile/migratory program. F. Median log_2_FC and gene-level heatmap for the inflammatory program. G. Collagen gel contraction assay in TGFβ-stimulated primary human cardiac fibroblasts treated with si*NREP*, si*TGFBI* or si*RBPJ*. Representative gel images at day 5. Quantification of percent contraction over time; mean ± SEM, n=3. Two-way ANOVA with Dunnett’s multiple comparisons test. H. Transwell migration assay in primary human cardiac fibroblasts treated with si*NREP*, si*TGFBI* or si*RBPJ*. Representative crystal violet–stained images and quantification of migration fold change relative to siCTRL; mean ± SEM, n=3. One-way ANOVA with Šídak’s multiple comparisons test.

## METHODS

*Note: References cited in the Methods section are numbered independently of the main manuscript and listed at the end of this section*.

### Immortalized Cardiac Fibroblast Culture

High glucose DMEM (Gibco 10566) supplemented with 10% FBS (SeraPrime F31016-500), 1X MEM NEAA (Gibco 11140050), 10 U/mL penicillin/streptomycin (Gibco 15140) and 1 mM sodium pyruvate (Gibco 11360070) was used to culture immortalized cardiac fibroblast cell lines. For routine maintenance of the cells, fibroblasts were passaged every 2-3 days and cultured in fresh medium. For TGFβ stimulation, fibroblasts were seeded at 80,000 cells per well in 6-well plates and cultured overnight. The following day, medium was replaced with serum-deprived medium (basal medium supplemented with 0.5% FBS instead of 10% FBS), and TGFβ (Fisher Scientific 100-21C) was added to a final concentration of 5 or 10 ng/mL, depending on the assay. Unless otherwise indicated, cells were stimulated for 48h.

### Cleavage Under Targets & Release Using Nuclease (CUT&RUN)

CUT&RUN was performed using the CUTANA ChIC/CUT&RUN Kit (EpiCypher 14-1048) to assess BRD4 occupancy in TGFβ stimulated BRD4-FLAG cardiac fibroblasts, RUNX1 binding in cardiac fibroblasts, and direct repression of targeted enhancers by the dHyperLbCas12a (dHLbCas12) system. Cells were seeded 24 h or stimulated 48 h before harvest, detached using TrypLE for collection, washed with 1× PBS, and resuspended in the CUT&RUN wash buffer. For each antibody, 500,000 (FLAG) or 1 × 10⁶ (RUNX1 and H3K9me3) fresh cells were prepared for two biological replicates. Samples were individually incubated with IgG (EpiCypher 13-0042, negative control), H3K4me3 (EpiCypher 13-0060, positive control), FLAG M2 (Sigma F1804-200ug), RUNX1 (ab220117), or H3K9me3 (Abcam ab8898) antibodies (1 µg per sample). Cells were first immobilized on concanavalin A beads and then incubated overnight at 4 °C with the appropriate antibody in antibody dilution buffer on a nutator. The following day, cells were washed twice with a cell permeabilization buffer and incubated with pAG-MNase for 1h at 4 °C. Calcium chloride was then added to a final concentration of 2 mM, and samples were incubated for an additional 2h at 4 °C to digest chromatin and release DNA. Fragmented DNA was purified and quantified using the Qubit 2.0 Fluorometer together with the Qubit dsDNA HS Assay Kit (Thermo Fisher Scientific). Paired-end Illumina sequencing libraries were prepared from CUT&RUN-enriched DNA using the NEBNext Ultra II DNA Library Prep Kit (New England Biolabs E7645) and sequenced on an Illumina NextSeq 1000/2000 using the XLEAPS-SBS Onboard Denature workflow. The nf-core/cutandrun pipeline (v.3.2.2)1 was used to perform primary analysis within the Nextflow bioinformatics workflow manager (v.25.04.6), using Conda for software dependency management. Reads were aligned to the mouse genome (GRCm39/mm39; GENCODE vM36 annotation) with a genome size parameter of 2,350,000,000 bp and default pipeline settings otherwise. The pipeline performs adapter trimming, read alignment and filtering, normalized coverage track generation, peak calling and annotation relative to gene features, consensus peak set creation, and quality control and version reporting.

### lentiMPRA Library Design and Construction

A lentivirus-based MPRA was used to assay the regulatory activity of 280 candidate enhancers identified by ATAC-seq and validated by CROP-seq. Each candidate enhancer was tiled into overlapping 140-bp candidate regulatory sequences (CRSs) with a 30-bp overlap, centered on the enhancer; candidate enhancers <1 kb were tiled into 2 CRSs and those >1 kb into 6 CRSs. Each CRS was flanked on both sides by a 15-bp adaptor, giving a final oligonucleotide length of 170 bp, and the resulting pool of 944 oligonucleotides was synthesized by GenScript (HiFi 92K).

The lentiMPRA library was constructed essentially as previously described2, with minor modifications. After pooled oligonucleotide synthesis, a two-round PCR was performed to pre-amplify the pool and to append barcodes and the additional sequences required for the lentiMPRA construct. Both rounds used Q5 High-Fidelity 2X Master Mix (New England Biolabs), with 20 and 15 cycles for the first-and second-round PCR, respectively. All primers used for library construction and sequencing-library preparation were synthesized by Integrated DNA Technologies (IDT). Purified PCR products were cloned by Gibson assembly into the pLS-SceI vector (Addgene 137724) linearized with AgeI-HF and SbfI-HF (New England Biolabs R3552 and R3642). The assembly was electroporated into 10-beta electrocompetent E. coli (New England Biolabs C3020) using a Gene Pulser Xcell electroporation system (Bio-Rad; 1-mm gap cuvette, 2 kV, 200 Ω, 25 µF, single 4.5-ms pulse). Bacteria from a single 10-cm plate were harvested and the plasmid library was purified over one column of a HiSpeed Plasmid Maxi Kit (QIAGEN).

### MRPA Plasmid Library Lentivirus Production

The plasmid library was co-transfected with the packaging plasmids psPAX2 (Addgene 12260) and pMD2.G (Addgene no. 12259) into HEK293T cells for lentivirus production. HEK293T cells (12 ×10⁶) were seeded in a 225-cm² flask and, the following day, transfected with 10 µg MPRA plasmid library, 6.5 µg psPAX2 and 3.5 µg pMD2.G using 60 µL FuGENE HD (Promega E2311). Eight hours after transfection the medium was replaced and supplemented with ViralBoost (ALSTEM VB100). Virus was harvested 3 days later, concentrated with Lenti-X Concentrator (Takara Bio #631231), resuspended in 600 µL DPBS and stored at 4 °C.

### lentiMRPA Transduction of Mouse Cardiac Fibroblasts

Mouse cardiac fibroblasts were maintained in high-glucose DMEM (Gibco 10566) supplemented with 10% FBS (Omega Scientific), 1 mM sodium pyruvate (Gibco 11360), 1× MEM-NEAA (Gibco 11140), and penicillin–streptomycin and were passaged at 10,000 cells cm⁻² using TrypLE (Gibco 12605). The following day, 33 µL of concentrated virus was added to each well of a 6-well plate in medium supplemented with 8 µg mL⁻¹ polybrene (Sigma TR-1003). The medium was replaced the next day. Two days later, cells were passaged into three 6-well plates at 3,000 cells cm⁻² (untreated) or 5,000 cells cm⁻² (treated). Untreated cells were maintained in the medium described above (10% FBS); treated cells were maintained in the same medium containing 0.5% FBS and supplemented with either 10 ng mL⁻¹ TGF-β1 (FisherScientific 100-21C) alone, or 10 ng mL⁻¹ TGF-β1 plus 0.5 µM JQ1 (MedChemExpress 1268524-70-4). After 48h, DNA and RNA were co-extracted using the AllPrep DNA/RNA Mini Kit (QIAGEN 80204), and sequencing libraries were prepared as previously described^2^. The CRS–barcode association library and the DNA/RNA barcode-count libraries were sequenced using the previously described parameters^2^ on an AVITI24 instrument (Element Biosciences) with AVITI 2×150 and AVITI 2×75 Cloudbreak Freestyle sequencing kits, respectively.

### MPRA Data Processing and Analysis

Raw reads from the CRS–barcode association and DNA/RNA barcode-count libraries were processed with MPRAsnakeflow (v0.5)^3^, using UMIs to collapse PCR duplicates. Downstream analysis was performed in Python (v3.13.7). Per-barcode activity was defined as the log2(RNA/DNA) count ratio; barcodes without a positive DNA-normalized ratio were discarded, and CRSs with fewer than three barcodes were excluded. For each comparison, conditions were merged on shared barcode–CRS pairs to retain only barcodes detected in both. The experiment was performed in two independent biological replicates. Within each replicate, every CRS was tested with a paired two-sided t-test comparing per-barcode log2(RNA/DNA) between conditions, pairing by barcode; P values were corrected across CRSs by the Benjamini–Hochberg procedure. Two comparisons were run: untreated versus TGFβ, and TGFβ versus TGFβ + JQ1. A CRS was called a hit at adjusted P < 0.05 with mean_delta > 0 (TGF-β1) or mean_delta < 0 (JQ1 co-treatment); a candidate enhancer was called active when at least one of its CRSs was a hit. The final set of responsive CRSs was the union of hits across the two replicates.

### CROP-seq Vector Modification

The CROP-seq vector (BSD-T2A-BFP) was modified from a previously published vector (Puromycin-T2A-mCherry)^4^. The original vector was first digested with BsiWI and MluI. The blasticidin resistance gene (BSD; gift from Dr. Alexander Marson; Addgene #252529) and T2A-BFP (Addgene #60955) were amplified using the primers outlined in the relevant “Tables” section. Amplified fragments were then ligated into the linearized vector using the Cold Fusion Cloning Kit (System Biosciences MC010B-1). BsmBI cut site within BFP (used for sgRNA cloning) was mutated using a site-directed mutagenesis kit (Agilent 200523) with the primers outlined in the relevant “Tables” section.

### CROP-seq Candidate Enhancer Selection

The initial set of candidate enhancers was scATAC-seq cell-enriched distal peaks in TAC^5^, either fibroblast-specific or non-fibroblast-specific. They were then filtered to satisfy the following criteria indicative of stress-activated fibroblast regulatory elements: (1) accessible in cardiac fibroblast scATAC-seq in the TAC model (accessibility > 0); (2) overlapping with broad H3K27ac peaks following in vitro TGFβ stimulation of cardiac fibroblasts^5^; (3) exhibiting PRO-seq signal following in vitro TGFβ stimulation (coverage > 0)^5^; (4) overlapping with BRD4 CUT&RUN peaks, called using SEACR (v.1.3) with the relaxed and normalized settings against the IgG control, identified in TGFβ-treated cardiac fibroblasts^6^; and (5) possessing syntenic human genomic regions, defined by lifting mm10 mouse candidate enhancer coordinates to hg38 human genome using liftOver function from R package rtracklayer (v.1.64.0) with UCSC mm10-to-hg38 chain file, retaining lifted regions ≥200 bp, assessing conserved genomic context by requiring lifted human regions to lie within 100 kb of orthologous human genes corresponding to genes located within 100 kb of the original mouse enhancer, collapsing all aligned human genomic intervals for each candidate enhancer into a single contiguous interval, and filtering resulting human genomic intervals (500 bp ≤ length ≤ 5,000 bp). Among the resulting candidate enhancers, 280 were subsequently selected based on satisfying one of the following criteria: (1) fibroblast-specific enhancers with strongly positive (coefficient > 5) or negative (coefficient < -5) fibroblast coefficients, calculated using the approach from Alexanian et al.^5^; (2) human syntenic regions overlapping peaks with increased accessibility in fibroblast from patients with AMI or chronic HF relative to healthy donors^7^; (3) human syntenic regions located within 150 kb of genes differentially expressed in AMI and chronic HF relative to healthy donors; (4) enhancer overlapping genes differentially expressed in TGFβ stimulation in both PRO-seq and bulk RNA-seq of immortalized mouse cardiac fibroblasts^5^.

### sgRNA Design and Pooled Lentivirus Library Preparation

Two sgRNAs were designed for each candidate enhancer using an automated CHOPCHOP (v.3, rev 2e9cfc9) workflow. Guide design was performed against the mouse reference genome mm10 to identify 20-nt SpCas9 guides with NGG PAM sites, using the Doench 2016 model to score guide activity. Candidate guides were restricted to a GC content of 20-80%, and no constraint was imposed on the 5’ dinucleotide preceding the guide sequence. Guides were subsequently filtered to retain guides with predicted efficiency >45%, no predicted off-target sites with up to two mismatches (MMO, MM1, MM2 > 0), and fewer than six predicted off-target sites with three mismatches (MM3 >= 6), and self-complementarity scores <2. Guides located outside of the ±500 bp window centered on the enhancer midpoint were excluded. The remaining guides were screened to identify the first pair in order of combined efficiency targeting distinct, non-overlapping positions within ±500 bp. Because 35 guides targeted regions shared between adjacent enhancers, the final library contained 525 unique targeting guides covering all 280 enhancers, plus 20 non-targeting controls (545 total). Primer3 was used to design PCR primers flanking each candidate enhancer target region with a search window of ±290 bp and a minimum guide-to-primer distance of 20 bp. Twenty previously validated non-targeting sgRNAs were included as controls^8^. The selected sgRNAs were cloned into a modified CROP-seq backbone (BSD-T2A-BFP) and used to produce a purified pooled lentivirus library (>10^8 TU/mL) by VectorBuilder (Chicago, USA).

### CROP-seq Lentivirus Transduction and TGFβ Stimulation of Mouse Cardiac Fibroblasts

Immortalized mouse cardiac fibroblast CRISPR interference (CRISPRi) line C1^5^ was seeded at a density of 10,500 cells cm^-2^. Cells were allowed to recover for 24 hours before being transduced with varying concentrations of lentivirus pool supplemented with 8ug/mL of polybrene (Sigma-Aldrich TR-1003-G) to ensure single sgRNA integration per cell following the manufacturer’s FACS-based lentivirus titration protocol (Millipore-Sigma). 72 hours post-transduction, flow cytometry analysis (BD Aria Fusion) with a portion of each transduction concentration was used to determine the percentage of BFP-positive cells. The dilution with the proper integration frequency was further cultured, subjected to blasticidin selection (4 ug/mL; ThermoFisher A1113903), and expanded with flow cytometry validation of percentage of BFP positive cells every three days. When the percentage of BFP-positive cells was greater than 90%, cells were plated, allowed to recover for 24 hours, and treated with 5 ng/mL TGFβ (FisherScientific 100-21C) in starvation medium (high glucose DMEM with 0.5% FBS, 1X Non-Essential Amino Acid (NEAA), 10 U/mL penicillin/streptomycin and 1 mM sodium pyruvate) for 48 hours. After the 48-hour treatment, cells were harvested for single cell library preparation. The number of cells harvested were seeded 1000 cell coverage per gRNA sequence.

### CROP-seq Single Cell Library Preparation and Data Analysis

Single cell transcriptome libraries were prepared by loading cells into the 10x Genomics Chromium X instrument (PN 1000690) according to manufacturer’s instructions. Further preparation was completed according to the user guide of the GEM-X Universal 3’ Gene Expression v4 kit (PN 1000691) and Dual Index Kit TT Set A (1000215). Final libraries were sequenced with NovaSeq X Plus (Illumina) with manufacture recommended cycling parameters (read 1: 28 cycles, i7 and i5: 10 cycles, and read 2: 90 cycles). Samples were demultiplexed, and fastq files were generated from raw sequencing data using Illumina DRAGEN BCL Convert (v.4.2.0). All samples were subsequently processed using the Cell Ranger pipeline (10x Genomics, v.9.0.1) as follows. Each sample was individually aligned to the mouse reference genome mm10 using cellranger count. To account for varying sequencing read depth, we ran cellranger aggr to normalize all samples to the mapped read depth of the least sequenced sample. Read-normalized aggregated values from cellranger aggr were used as initial input using the R package Seurat (v.5.3.0) using the functions Read10X and CreateSeuratObject with default parameters. Each sample used in aggregation was identified by run and assigned a unique name in metadata as ‘gem.group’. Quality control filtering included removal of outliers due to the number of features/genes (nFeature_RNA >= 600 and nFeature_RNA <= 3,000), UMI counts (>= 900 and <= 100,000), mitochondrial percentage (<= 6%), and ribosomal percentage (<= 30%). SCT normalization was then performed. Principal component analysis and batch correction using Harmony were then performed using split.by = “gem.group”. Clustering was then run using the functions RunUMAP, FindNeighbors, and FindClusters, and the output UMAP graphs were generated by DimPlot. Differential gene expression of each enhancer knockdown target and non-targeting control cells was performed using a Wilcoxon rank-sum test implemented in wilcoxauc (Presto, v.1.0.0) for e2g and proximal genes.

### sgRNA Amplicon Library Generation

sgRNA amplicon library generation was completed as previously described^9^. In short, 13 ng of cDNA was taken after the cDNA purification step in the 10x Genomics’s GEM X Universal 3’ Expression single cell library preparation and subjected to three rounds of nested PCRs using the KAPA HiFi polymerase (Roche KM2605). Primers for the nested PCRs are listed below and were used at a final concentration of 0.3 µM. The amplicon libraries were then purified with SPRISelect Beads (Fisher Scientific NC0406406) and assessed with the Bioanalyzer HS DNA kit (Agilent 5067) prior to pooling. The pooled amplicon library was sequenced on the Illumina NextSeq 2000 (Illumina) using the same cycling parameters as described above for the endogenous scRNA-seq libraries. Detailed sequences for primers used are outlined in the relevant “Tables” section.

### Amplicon Library Analysis

Amplicon libraries were demultiplexed, and fastq files were generated from raw sequencing data using Illumina DRAGEN BCL Convert. Cell Ranger count was used to generate BAM files corresponding to each sample’s endogenous transcriptome. Guide assignments were subsequently determined from the bam files as previously described^9^, using a whitelist of all 585 sgRNA sequences in our library to generate a table of cells, barcodes (sgRNA sequences), read counts, and UMI counts. Cell barcodes with zero read counts were removed, followed by exclusion of barcodes that could not be confidently extracted from the sequencing reads. Remaining barcode sequences were matched to the reference sgRNA library to assign guide identities. To ensure high-confidence perturbation assignments, cells containing more than one unique targeting (i.e., non-control) sgRNA were excluded from downstream analyses, whereas multiple non-targeting control sgRNAs were retained as a single control category. The filtered sgRNA assignments were then matched to the corresponding cell barcodes in the Seurat object described above. We then assigned sgRNA sequence and enhancer knockdown target to a specific cell as a metadata value. All cells were associated with one of the 268 candidate enhancers (after preprocessing, we obtained 268 enhancers from the initial 280), and all 20 non-targeting control sequences were considered together as control cells.

### Gene program annotation

A joint supervised–unsupervised non-negative matrix factorization framework, ALPINE^10^ (Adaptive Layering of Phenotypic and Integrative Noise Extraction), was used to identify perturbation-associated gene programs and their marker genes. An ALPINE model was trained using sgRNA_target as the perturbation covariate (n_components = 54, n_covariate_components = 15, alpha_W = 0, lam = 825). The optimal number of covariate components was determined empirically by training ALPINE models with n_covariate_components ranging from 0 to 20 and evaluating the reconstruction loss. Fifteen covariate components were selected since the loss curve exhibited diminishing returns beyond this point. The input consisted of SCTransform-normalized expression values for the 5,000 highly variable genes from CROP-seq fibroblasts with a fibroblast cell-state score above the 70th percentile of non-targeting control cells. The fitted model was used to generate cell embeddings and the decomposed matrices W, H, and B, representing gene-by-program loadings, program activities across cells, and perturbation-by-program effects, respectively. These outputs were subsequently used to identify perturbation-associated gene programs, cluster genes according to their program activities, and define program-specific marker genes for downstream biological interpretation. The 15 gene programs identified by ALPINE were annotated based on the top-ranked genes in each program’s gene loading vector (matrix W). For each program, the 25 highest-loading genes were examined for known biological functions, and pathway enrichment was performed using Reactome. Programs were assigned to ten biological categories based on the convergence of top gene identities and pathway enrichment: contractile myofibroblast (P5, P6, P10), ECM deposition (P9), migration and paracrine signaling (P7), myofibroblast proliferation (P13, P14), metabolic activation (P2), chromatin remodeling and stress (P3), ECM sensing and turnover (P15), profibrotic BMP-redox (P12), TGFβ autocrine (P4, P5), and quiescence (P1). Category assignments were based on manual inspection of gene function and pathway enrichment results and are not outputs of the ALPINE algorithm itself.

### Scoring Candidate Enhancer Knockout Effects

An elastic net–regularized generalized linear model was trained using the glmnet engine implemented through the tidymodels framework (v.1.2.0) to score the healthy-to-diseased fibroblast cell state following CROP-seq perturbations. Training data consisted of SCTransform-normalized scRNA-seq expression values. Separate models were trained using human donor versus chronic HF fibroblasts and mouse sham versus TAC fibroblasts as described above. For each model, candidate predictors were restricted to highly variable genes identified in both the training scRNA-seq and CROP-seq datasets using the FindVariableFeatures function (nfeatures = 5000) in the Seurat package. For the human model, mouse genes were mapped to their corresponding human orthologs using biomaRt (v.2.60.0). Candidate predictors were restricted to one-to-one mouse–human orthologs to enable cross-species application of the model. Cells were randomly divided into training (70%) and test (30%) sets using stratified sampling. Predictors with zero variance and highly correlated features (Pearson correlation coefficient >0.9) were removed before model fitting. Hyperparameters controlling the regularization penalty (λ) and elastic net mixing parameter (α) were optimized by grid search with 10-fold stratified cross-validation. Model performance was evaluated on held-out test cells using the root mean squared error (RMSE) and coefficient of determination (R²). Before the trained models could be applied to the CROP-seq dataset to make predictions, CROP-seq expression profiles were restricted to the selected predictor genes, and expression values were adjusted by a global library-size scaling factor to account for differences in sequencing depth between datasets. To score the CROP-seq cells using the human model, mouse genes were mapped to their corresponding human orthologs as above. The optimized models were subsequently applied to SCTransform-normalized CROP-seq expression profiles. Linear model predictions were transformed to probabilities using the logistic function to generate a continuous fibroblast cell-state score for each perturbed cell, with higher scores indicating greater similarity to the healthy fibroblast transcriptional state.

### CRISPRi knockdown of transcription factors

gRNAs targeting each transcription factor (TF) were selected from the mCRISPRi-v2 sgRNA library^11^ (sequences listed in Supplementary Table). For each TF, two gRNA oligo pairs, each with 5′ (Sense: 5′-TTG; Antisense: 5′-TTAGCTCTTAAAC) and 3′ (Sense: 3′-GTTTAAGAGC; Antisense: CAACAAG) overhangs, were annealed and phosphorylated using T4 Kinase (New England Biolabs) and sub-cloned into BstXI-and BlpI-double-digested pU6-sgRNAEF1Alpha-HygT2A-BFP (modified from pU6-sgRNAEF1Alpha-puroT2A-BFP, gift from Dr. Jonathan Weissman, Addgene #60955, by replacing the puromycin resistance gene with a hygromycin resistance gene) using T4 ligase-mediated ligation. Constructs were sequence-verified (Quintara Bio, Berkeley, CA, USA). To generate lentiviral particles, 1 × 10⁶ HEK293T cells were seeded on a 100 mm plate one day prior to transfection and cultured in 10 mL fibroblast medium. On the day of transfection, medium was replenished with 10 mL fresh medium. 5 µg of the gRNA lentiviral vector (2.5 µg of each gRNA vector) was co-transfected with 2.5 µg of envelope plasmid pMD2.G (Addgene #12258) and 2.5 µg of packaging plasmid psPAX2 (Addgene #12260) into HEK293T cells using 59 µL of FuGENE HD transfection reagent (Promega E2312), following the manufacturer’s instructions. Supernatant was collected 48 h after transfection. All supernatants were filtered through 0.45 µm PVDF syringe filters (Thermo Fisher Scientific 723-9945) to remove cell debris. The previously generated CRISPRi fibroblast clone (C1)^5^, stably expressing KRAB-dCas9-mCherry (pHR-SFFV-KRAB-dCas9-mCherry; gift from Dr. Jonathan Weissman, Addgene #60954), was transduced with 2 mL of each of the two gRNA-containing lentiviral supernatants targeting each TF. Cells harboring the targeting gRNAs were selected by hygromycin treatment at 100 µg/mL for 7 days to generate a pure polyclonal population.

### Bulk ATAC-seq

Bulk ATAC-seq was performed using the Omni-ATAC protocol^12^ with three biological replicates per condition (unstimulated and TGFβ-stimulated). Fibroblasts were seeded at 1 × 10⁵ cells per well in 6-well plates 24 h before stimulation and treated with 5 ng/mL TGFβ. After 48 h, cells were detached using TrypLE, washed once with 1× PBS, and counted. For each sample, 60,000 cells were pelleted at 500×*g* for 5 min at 4 °C. Nuclei were isolated by resuspending cell pellets in cold lysis buffer (10 mM Tris-HCl, pH 7.4, 10 mM NaCl, 3 mM MgCl₂, 0.1% Tween-20, 0.1% NP-40, 0.01% digitonin, and 1% BSA) and incubating for 10 min at 4 °C. Nuclei were washed in ATAC-seq wash buffer (10 mM Tris-HCl, pH 7.5, 10 mM NaCl, 2 mM MgCl₂, and 0.1% Tween-20) and pelleted at 500×*g* for 10 min at 4 °C. Purified nuclei were then resuspended in 50 µL transposition reaction mix containing 1× TD buffer (Diagenode C01019043), TDE1 Tn5 transposase (Diagenode C01070013) with 100 nM final concentration, 0.01% digitonin, and 0.1% Tween-20. Transposition reactions were incubated at 37 °C for 30 min with shaking at 1,000 rpm. Transposed DNA was purified using DNA Clean & Concentrator-5 columns (Zymo Research D4014) and amplified with indexed Ad1/Ad2 adapters (0.25 µM each) using the NEBNext Ultra II Q5 2× Master Mix (New England Biolabs M0544). Libraries were initially amplified for five PCR cycles, and the number of additional cycles was determined from the post-barcoding library concentration measured using the Qubit dsDNA HS Assay Kit (Thermo Fisher Scientific Q33231) according to the formula: N = −5.7 × log₁₀[ng/µL] + 6.7. Amplified libraries were purified, quantified using the Qubit dsDNA HS Assay Kit (Thermo Fisher Scientific Q33231), and assessed for fragment size distribution using an Agilent Bioanalyzer. Paired-end libraries were sequenced on an Illumina NextSeq 1000/2000 with a target depth of 20–50 million read pairs per sample. Bulk ATAC-seq raw reads were aligned to the GRCm38 (mm10) reference genome using nf-core/atacseq (v.2.1.2)^13^. A consensus peak set was generated, and peak-level read counts were obtained using featureCounts from the consensus_peaks.mRp.clN.featureCounts.txt output. For differential accessibility analysis, peaks with no accessibility across samples were removed. Filtered count data were transformed using the voom method in limma (v.3.60.0), with quantile normalization, to generate log2-counts per million (log2-CPM) values and model the mean–variance relationship. Differential accessibility was assessed using linear models with empirical Bayes moderation implemented in limma (v.3.60.0).

### Bulk RNA sequencing

Cells were collected and lysed directly in DNA/RNA Shield (Zymo Research, R1100) and submitted to Plasmidsaurus for bulk RNA sequencing. Bulk RNA-seq raw reads were aligned to the GRCm38 (mm10) genome using nf-core/rnaseq (v.3.12.0). Gene-level read counts were obtained from the Salmon.merged.gene_counts.rds output. For differential expression analysis, genes with maximum expression <0.5 counts per million (CPM) across samples were removed. Filtered count data were transformed using the limma voom method from the Bioconductor package edgeR (v.4.2.0) to generate log2-CPM expression values and model the mean-variance relationship. Differential expression was assessed using linear models with empirical Bayes moderation using Limma (v.3.60.0). For visualization of siRNA knockdown effects of the 10 effector genes (SERPINE1, ADAM12, BGN, COL5A1, SOCS5, LRRC32, PSD3, EDN1, MICAL2, AEBP1), differential expression was performed with DESeq2 (v.1.44.0), comparing each siRNA condition against the NCTRL2_1 control (n = 4 biological replicates per condition). Gene-set–level shifts were assessed with a two-sided Mann–Whitney U test comparing per-gene log₂FC values against zero, using two GO annotations (ECM Organization, GO:0045229, 84 genes; Inflammatory Response, GO:0006954, 163 genes) and two curated fibroblast-specific sets (32 Contractile genes and 25 Migration genes). Specifically for the analysis of the effects of knocking down *NREP, TGFBI* and *RBPJ*, gene sets were taken unmodified from MSigDB v7.0, combining the Hallmark collection with the GO biological process collection, and terms were selected by name for the two processes of interest before any result was inspected: seven proliferation terms (Hallmark E2F targets, G2M checkpoint, MYC targets V1 and mitotic spindle; GO mitotic cell cycle, DNA replication and chromosome segregation) and eight migration terms (Hallmark epithelial–mesenchymal transition; GO locomotion, cell motility, amoeboidal-type cell migration, tissue migration, smooth-muscle cell migration, cell–matrix adhesion and negative regulation of cell motility). Each set was intersected with the measured genes, giving 62–1,297 genes per set. For every set and knockdown, the log₂ fold changes of the set genes were compared with those of all other measured genes by a two-sided Mann–Whitney test, a competitive and threshold-free test that uses the full ranking rather than a differential-expression cut-off; P values were adjusted by Benjamini–Hochberg across the 45 set-by-knockdown comparisons. Effect size is reported as the mean log₂ fold change of the set genes, and is interpreted alongside the adjusted P value because the largest sets (>1,000 genes) reach significance on mean shifts below 0.05, which are not biologically meaningful. Per-set statistics, set sizes and the fraction of set genes individually reaching FDR < 0.05 are provided in the accompanying source data.

### *Meox1^flox/flox^* mouse line generation and validation

*Meox1^flox/flox^* mice were generated and purchased from GemPharmatech (Nanjing, China). To validate the conditional knockout allele, *Meox1^flox/flox^* animals were crossed with a *BetaActin^Cre/+^* animal (JAX: 033984)^14^ to generate whole body knock out animals. Animals were genotyped at 2 weeks of age (primer outlined in relevant “Tables” section), and the phenotype of the full body knock out animals was observed at 8 weeks of age in comparison to wild type and heterozygous knock out littermates. Timed matings of *Meox1^flox/flox^* animals and a *BetaActin^Cre/+^* animal were set up, and vaginal plug checks were conducted in the morning. Ultrasound validation of pregnancy was conducted at E6.5 (Vevo 3100 High Resolution Imaging System, FujiFilm VisualSonics). E10.5 embryos were collected by euthanizing the pregnant female and collecting embryos in ice-cold PBS supplemented with FBS (SeraPrime F31016-500) and stored on ice. During dissection, the yolk sac was isolated and DNA for genotyping was extracted from it using 50 µL of QuickExtract DNA Extraction Solution (Biosearch Technologies QE09050). E10.5 embryos were then mechanically digested in QIAzol (Qiagen 79306) and spun down to remove debris. RNA was then extracted using the Qiagen RNeasy Mini Kit (Qiagen 74104) and converted into cDNA using the iScript cDNA Synthesis Kit (Bio-Rad 1708841). Real-time quantitative PCR (RT qPCR) was conducted using a Taqman probe to measure *Meox1* transcript abundance (ThermoFisher 4304437; Mm00440285_m1).

### Transverse Aortic Constriction Surgery on Activated Fibroblast-Specific Meox1 KO Mice

Activated fibroblast-specific Meox1knock out mice were generated by crossing a Postn^mcm/+^ animal (JAX: 029645)^15^ with Meox1^flox/flox^ animals mentioned above. Postn^mcm/+^; Meox1^flox/flox^ and Postn^+/+^; Meox1^flox/flox^ littermates were subjected to transverse aortic constriction (TAC) or sham surgery between 8-10 weeks old. During surgery, body temperature was maintained at 37°C using a temperature-controlled small-animal surgical table after anesthesia with isoflurane (Harvard Apparatus). Mice were then subjected to thoracotomy. As previously described for TAC^16^, using 7-0 silk sutures and a 25-gauge needle, the aortic arch was constricted between the left common carotid and brachiocephalic arteries. For sham surgeries, thoracotomy was performed and the aorta was surgically exposed without further intervention. 100 mg/kg of tamoxifen (Sigma-Aldrich T5648) dissolved in corn oil (Sigma-Aldrich C8267) was injected intraperitoneally daily for 10 days starting 7 days post-surgery. Tamoxifen regimen was then changed to every other day at the end of the continuous treatment until the mice were harvested.

### Echocardiography Assessment of Cardiac Function

Echocardiography to evaluate cardiac function was performed prior to surgery, 4 weeks post-surgery, and 6 weeks post-surgery. As previously described^16^, mice were anesthetized with 1% isoflurane, and images were acquired using the Vevo 3100 (FujiFilm VisualSonics) and the MX550S probe. Briefly, long-axis and short-axis views were recorded in B-Mode and M-Mode, with high-frame-rate recordings obtained by electrocardiogram-gated kilohertz visualization (EKV) and from parasternal long-axis B-mode loops for speckle-tracking strain analysis. LV dimensions and ejection fraction were calculated from B-Mode and M-Mode, and global longitudinal strain (GLS)^17^ quantified using the VevoStrain speckle-tracking software package (FujiFilm VisualSonics). All analysis was performed blinded, and all reported parameters were averaged across multiple recordings. Animals that died before the 6-week endpoint were excluded from echocardiographic analyses.

### Collection and Fixation of Hearts

Hearts were perfused with cold PBS through the left ventricle after puncturing the portal vein and collected 6 weeks post-TAC and sham surgeries. The hearts were fixed overnight in 4% paraformaldehyde (PFA, Santa Cruz Biotechnology sc-281692) in 4°C. Samples were then washed with PBS three times, each for 5 minutes and stored in 30% sucrose for 24 hours at 4°C. The hearts were then embedded in OCT (Fisher Scientific 23-730-571) and cryosectioned using the Cryostar NX50. Cryosections were stored in -80°C until downstream usage.

### Picrosirius Red Staining and Quantification

For staining, each section was post-fixed with 4% PFA for 15 minutes at room temperature (RT). The sections were then washed with PBS 3 times for 5 minutes each. Sections were then stored in Bouin’s Fixative Substitute (Fisher Scientific NC9772216) overnight at RT. After a wash with DI water, the sections were stained with 0.1% Fast Green FCF (Thermo Fisher Scientific A16520.14) dissolved in 1% acetic acid for 3 minutes. The sections were rinsed off with DI water, washed with 1% acetic acid for 1 minute, and again rinsed with DI water. Sections were stained with picrosirius red solution (Polysciences 24901B-500) for 1.5 minutes. The sections were then washed twice with 1N HCl for 1 min each, once with 70% ethanol for 30 seconds, once with 100% ethanol for 30 seconds, and then stored in xylene for 5 minutes. The sections were then air-dried and mounted with ProLong Gold Antifade Mountant (ThermoFisher P36930). Sections were imaged using Keyence BZX810. Picrosirius red-positive area was quantified using Fiji (ImageJ) and expressed as a percentage of the total region of interest.

### Langendorff Perfusion and Single Cell Isolation

6 weeks post-surgery, the hearts of the mice were harvested, and non-cardiomyocytes (non-CMs) were isolated through Langendorff perfusion as previously described with minor changes^18^. To elaborate, mice were properly anesthetized and the heart was isolated after thoracotomy. The heart was then cannulated and perfused using a Langendorff apparatus (Radnoti 120108EZ) for 5-10 minutes at 37°C with perfusion buffer with final concentrations of 120.4 mM NaCl, 14.7 mM KCl, 0.6 mM KH2PO4, 0.6 mM Na2HPO4, 1.2 mM MgSO4, 10 mM Na-HEPES, 4.6 mM NaHCO3, 30 mM taurine, 10 mM 2,3-butanedione monoxime, and 5.5 mM glucose, pH 7.0. The heart was then digested with 300 units/mL collagenase II (Worthington Biochemical LS004177) and 50 μM CaCl_2_ dissolved in perfusion buffer for 10 mins at 37°C. The atria and greater vessels were removed following digestion, and the ventricles were teased apart into small pieces in perfusion buffer with 10% fetal bovine serum (FBS; stop buffer) at 37°C. After disrupting the tissue, the cell suspension was passed through a 250 μm strainer into a 50 mL falcon tube. The tube was then centrifuged at 30xg for 3 minutes at 4°C. The pellet containing the CMs was collected for downstream applications. The supernatant containing non-CMs was then collected and further filtered through a 70 μm cell strainer into a 15 mL tube. The 15 mL tube was then centrifuged at 400xg for 3 minutes at 4°C. The supernatant was then removed and then pellet was resuspended in 15mL of cold PBS for a wash. The cells again were centrifuged at 400xg for 3 minutes at 4°C, and the supernatant, again, was removed. The cells were then resuspended in 300 μL of cold PBS supplemented with 5% FBS and placed on ice for downstream applications.

### Sorting of Non-Cardiomyocyte Cells

Following isolation of non-CMs from Langendorff perfusion, a small fraction of the cells was taken for cell counting. The cells were transferred into a 1.5 mL microcentrifuge tube and spun down at 500xg for 5 minutes at 4°C. The supernatant was removed and the cells were resuspended in 50 μL of live/dead stain, NIR-APC-Cy7 (ThermoFisher L34975), per 1 million cells for 20 minutes on ice in the dark. 1mL of cold, fluorescent-activated cell sorting (FACS) buffer (PBS with 5% FBS) was added to quench the staining, and the cells were then centrifuged at 500xg for 5 minutes at 4°C. After the supernatant was removed, the cell pellet was resuspended in 100-200uL of antibody mix (FACS buffer with 1:50 CD16/32 (FC Block) (Cytek 70-0161-U500), 1:100 CD45-PerCP-Cy5.5 (Biolegend 103132), 1:100 CD31-PE-Cy7 (Biolegend 102418), 1:50 MEFSK4-APC (Miltenyi Biotec 130-120-802)) for 40 minutes on ice in the dark. 1 mL of cold FACS buffer was used to quench the antibody staining and cells were spun at 500xg for 5 minutes at 4°C. Two more washes were done after the supernatant was removed from the first wash. Cells were resuspended in 300-500 μL of cold FACs buffer and filtered through a 35 μm filter into FACS tubes. Cardiac fibroblasts were then sorted on the BD Aria Fusion gating for MEFSK4^+^, CD45^-^, and CD31^-^cells. Immune cells were sorted using the MEFSK4^-^, CD45^+^, and CD31^-^ gate. Endothelial cells were sorted using the MEFSK4^-^, CD45^-^, and CD31^+^ gate.

### Single-Cell Transcriptome Library Preparation, Sequencing, and Processing of non-cardiomyocyte cells

Non-cardiomyocytes obtained post Langendorff perfusion in *Meox1*^flox/flox^ and *Postn*^MCM^; *Meox1*^flox/flox^ TAC mice were fixed using the 10x Genomics’ Single Cell Fixed RNA Sample Preparation Kit (PN 1000414) according to manufacturer’s instruction and stored at -80 °C until library preparation. After thawing the frozen cells on ice, single-cell libraries were generated using the 10x Genomics Chromium X according to manufacturer recommendation from the Chromium Next GEM Flex Gene Expression kit (PN 1000497) User Guide. Additional components to generate the library are the Chromium Next GEM Chip Q Single Cell Kit (PN 1000422) and the Dual Index Kit TS Set A (PN 1000251). Final libraries were then sequenced with NovaSeq X Plus (Illumina). Raw sequencing reads were demultiplexed and aligned to the mm39 reference genome using Cell Ranger with the 10x Chromium FLEX probe set. Downstream analysis was performed in R using Seurat (v5.5.1). Counts were normalized with SCTransform, regressing out nFeature_RNA and mitochondrial gene percentage (percent.mt). Principal component analysis was computed on the SCT-normalized matrix (50 PCs), and dimensionality reduction (UMAP), nearest-neighbor graph construction, and unsupervised clustering (Louvain algorithm, resolution 0.1) were performed on the first 40 principal components. Clusters were annotated by inspection of canonical lineage markers. Differential expression analysis was performed within each broad cell type using Seurat’s FindMarkers (Wilcoxon rank-sum test) on log-normalized RNA counts, with thresholds |log₂FC| > 0.25, min.pct = 0.10, and adjusted p-value < 0.05 (Bonferroni correction). Pathway enrichment was performed with enrichR against the GO Biological Process 2023 database.

### RNAscope to Detect Meox1 Expression

Cryosections from sham and TAC hearts were removed from -80°C storage and hybridization of the *Meox1* and *Tnnt2* RNAscope probes were done with modifications from the RNAscope 2.5 HD Detection Kit (RED) Quick Guide and the RNAscope 4-plex Ancillary Kit for Multiplex Fluorescent Reagent Kit v2 Technical Note. Slides were baked at 60°C for 1 hour. Using the reagents in the Multiplex Fluorescent Reagent Kit v2 (ACD 323100), 5-8 drops of Hydrogen Peroxide were added to the sections and incubated at room temperature (RT) for 10 minutes. Slides were washed in distilled water twice. Antigen retrieval was completed by submerging the slides into boiling 1X Target Retrieval solution that was previously prepared from the 10X stock and was maintained at a mild boil for 5 minutes. After the 5 minutes, slides were washed with distilled water twice and rinsed with 100% EtOH before having excess liquid wiped off (taking care not to disrupt the tissue). A hydrophobic barrier was drawn around the sections before ∼5 drops of Protease Plus was added. The slides were then incubated in 40°C for 30 minutes followed by two washes with distilled water. Mouse *Meox1* C2 (ACD 530641-C2) and *Tnnt2* C4 (ACD 418681-C4) probes were added to RNAscope Probe Diluent and warmed at 40°C for 10 minutes before adding to dried slides for incubation at 40°C overnight. After the overnight incubation, slides were washed in freshly prepared 1X Wash Buffer at RT for 2 minutes. 1X Wash Buffer was prepared by diluting the 50X Wash Buffer (prewarmed to 40°C for 10-20 minutes) with distilled water. Hybridization of Amp 1 and 2 were done consecutively following the same procedure of dropping solution onto the slide and incubating at 40°C for 30 minutes and then washing twice with 1X Wash Buffer. Amp 3 was then added to cover the slide followed by a 15-minute incubation at 40°C before washing twice with 1X Wash Buffer. Development of HRP-C2 and HRP-C4 signals was done consecutively with the following protocol. 4-6 drops of the respective RNAscope Multiplex FL v2 HRP-C2 or HRP-C4 was added to the slide. Slides were incubated at 40°C for 15 minutes and then washed twice at RT with 1X Wash Buffer. An opal fluorophore (diluted 1:500 in RNAscope Multiplex TSA) were added to the slides, followed by a 30-minute incubation at 40°C. Opal 520 was used for *Meox1* C2 and Opal 690 was used for *Tnnt2* C4. After the 30-minute incubation, wash slides twice with 1X Wash Buffer at RT and dry slides before adding 4-6 drops of RNAscope Multiplex FL v2 HRP blocker. Slides were again placed in a 40°C oven and incubated for 15 minutes followed by two washes with 1X Wash Buffer at RT. After developing the HRP signals, slides were patted dry and ∼4 drops of DAPI were added and incubated for 30 seconds at RT. Slides were lightly tapped on the side to remove DAPI and glass coverslips were mounted on with Prolong Gode antifade mounting media. Slides were dried in the dark in overnight and stored at 4°C before imaging.

### cNMF analysis for in vivo gene programs

Raw count matrices from the *in vivo* fibroblast Seurat object were exported to H5AD format using the R anndata package, retaining the unnormalized count slot. Consensus non-negative matrix factorization (cNMF)^19^ was run across a range of K = 5–10 components with 100 iterations per replicate (n = 10,000 total NMF solutions). Outlier NMF solutions were identified and removed using a local density threshold of 0.01; 2,893 of 10,000 spectra (29%) were excluded prior to consensus clustering. K selection was guided jointly by program stability (silhouette score across replicates) and reconstruction error: stability was highest and most consistent at K = 5–6 (>0.88), dropped sharply at K = 7–8, then recovered at K = 10, while reconstruction error decreased monotonically. K = 10 was selected to balance resolution and stability, supported by the well-separated block-diagonal structure of the consensus distance matrix at this K. Cell usage scores (cells × programs) and gene spectra scores (programs × genes) were loaded back into R and integrated with the Seurat metadata. Three programs (cNMF_8, cNMF_9, cNMF_10) were identified as ambient/contamination signatures based on their top-weighted genes (endothelial, T/NK cell, and neutrophil markers, respectively) and were excluded from downstream analyses, yielding seven fibroblast-intrinsic programs. Program identity was characterized using the top 25 genes by spectra score. Differential program usage between conditions was assessed per cell using a two-sided Wilcoxon rank-sum test with Benjamini–Hochberg correction. Gene set enrichment within each program was computed as the mean z-scored spectra score across genes of interest. Similarity between in vivo cNMF programs and in vitro programs was quantified by averaging in vivo spectra scores over the top 50 genes of each in vitro program, followed by row-wise z-scoring of the resulting similarity matrix. All visualizations were produced in R using ggplot2, pheatmap, and ComplexHeatmap.

### Culturing of human primary cardiac fibroblasts and siRNA transfection

Human primary cardiac fibroblasts (Lonza CC-2904) were cultured in FibroGRO Complete Medium (Millipore Sigma SCMF001) with 2% FBS at 37°C in a humidified incubator with 5% CO₂. For TGFβ stimulation, cells were cultured in FibroGRO Basal Medium with 0.5% FBS and addition of TGFβ (10 ng/mL; PeproTech, 100-21C). For serum starvation only, cells were cultured in FibroGRO Basal Medium with 0.5% FBS. Fibroblasts were used at passage 6–8, and passage number was kept consistent within each assay type. 100,000 human primary cardiac fibroblasts were seeded in a six-well plate the day before transfection. Wells to be stimulated with TGFβ on the day of transfection were switched to FibroGRO Basal Medium with 0.5% FBS, while unstimulated wells were switched to FibroGRO Basal Medium with 2% FBS. Cells were transfected with a final concentration of 15 nM siRNA (Dharmacon ON-TARGETplus human siRNA SMARTPool, Horizon Discovery Biosciences; details provided in the Supplementary Tables) using 7 µL of Lipofectamine RNAiMAX transfection reagent (Thermo Fisher Scientific 13778150) per well, according to the manufacturer’s instructions. Then, 24 h after transfection, appropriate wells were treated with TGFβ or maintained in serum starvation. Cells were collected 48 h after stimulation.

### Collagen Gel Contraction Assay

Tissue culture-treated 24-well plates were pre-coated with 2% BSA in PBS. Compressible collagen matrices (1.25%) were prepared in the wells using PureCol EZ Gel Solution (Advanced BioMatrix #5074) and polymerized at 37 °C for 1.5h. Fibroblasts suspended in serum-supplemented medium were seeded onto the collagen gels at a density of 50,000 cells per gel and cultured for 2h. To initiate gel contraction, the medium in unstimulated wells was replaced with fresh serum-supplemented medium, whereas the medium in TGF-β-stimulated wells was replaced with serum-deprived medium containing 10 ng/mL TGFβ. Gels were gently released from the well walls, and cells were cultured for up to 96 h. Brightfield images were acquired every 24h. Gel area was quantified using FIJI (ImageJ), and contraction was expressed as the percentage of contraction in gel area relative to the initial gel area.

### Transwell cell migration assay

Cell migration was assessed using 24-well Transwell inserts with 8-µm-pore polycarbonate membranes (Corning 07-200-174). Fibroblasts (10,000 cells in 100 µL) suspended in either serum-supplemented medium or serum-deprived medium containing TGFβ were seeded into the upper chamber, and 600 µL of serum-supplemented medium was added to the lower chamber. After incubation for 24 h at 37°C, medium was removed, and inserts were washed three times with 1× PBS. Cells remaining on the upper surface of the membrane were removed with a moistened cotton swab. Migrated cells on the lower surface were fixed with 4% paraformaldehyde for 20 min and stained with 0.2% crystal violet (Sigma-Aldrich, 61135) for 20 min. Membranes were washed with 1× PBS until excess crystal violet was completely removed. Migrated cells were imaged in PBS using a Keyence BZ-X810 microscope. Five randomly selected fields of view (10× objective) from top, bottom, left, right, and center of each membrane were imaged per membrane, and the mean number of migrated cells per field was quantified using FIJI (ImageJ).

### α-Smooth Muscle Actin Immunofluorescence Staining and Quantification

Mouse cardiac fibroblasts were seeded onto poly-D-lysine-coated glass coverslips in 6-well plates. After 24h, cells were either stimulated with 10 ng/mL TGFβ or left untreated and cultured for another 48h. Cells were fixed with 4% paraformaldehyde for 20 min and washed three times with 1× PBS for 5 min each. Samples were permeabilized with 0.1% Triton X-100 (Sigma-Aldrich) for 30 min and incubated overnight at 4 °C with a primary antibody against α-smooth muscle actin (αSMA; Abcam ab7817; 1:300 dilution in 5% BSA in PBS). After washing with 1× PBS, cells were incubated with an Alexa Fluor-conjugated secondary antibody Alexa 488 (Thermo Fisher Scientific A-11029) for 1h in the dark, followed by DAPI staining (1 µg/mL) for 10 min. Coverslips were mounted using ProLong Diamond Antifade Mountant (Thermo Fisher Scientific P36930) and cured overnight. Images were acquired using a Keyence BZ-X810 microscope with identical acquisition settings for all samples. αSMA expression was quantified using FIJI (ImageJ). DAPI images were thresholded and analyzed using the particle analysis function to identify nuclei, and objects below a predefined area threshold (0.01 pixel²) were excluded from analysis. Total αSMA fluorescence was quantified as the integrated fluorescence intensity across the entire field of view and normalized to the number of retained nuclei to obtain the average αSMA fluorescence intensity per nucleus. Eight randomly selected fields of view were analyzed for each well.

### Isolation and culture of mouse BMDMs

BMDMs were isolated from 6–10 week-old male or female C57BL/6J (B6J) mice. After mice were euthanized, tibias and femurs were removed, cannulated with a 25-gauge needle and flushed with sterile DMEM (Thermo Fisher Scientific SH30022FS) containing 10% bovine growth serum (BGS; Thermo Fisher Scientific SH3054103) and 1% penicillin/streptomycin (Gibco 15140) to isolate whole-bone-marrow cells. Then, 4 × 10^5^ cells were plated and cultured on a 10 cm uncoated Petri plate in DMEM containing 10% bovine growth serum, 1% penicillin–streptomycin and 50 ng/ml macrophage colony stimulating factor (MCSF; ProSpecBio CYT-439). A 50% medium refresh was performed after 3 days and cells were then split onto assay plates after reaching 70% confluency (days 7–8 after plating).

### TGFβ stimulated fibroblasts for conditioned medium experiments on BMDMs and gene expression analysis

Fibroblasts were cultured and stimulated as described in “Immortalized Cardiac Fibroblast Culture and TGFβ Stimulation.” After 48 h incubation, the medium was collected, centrifuged at 500 × g for 10 min at 4°C, and passed through a 0.2 µm PES filter (Pall Life Sciences, 4612) to remove cell debris. Passage 1 BMDMs, grown and differentiated as described above, were plated overnight on six-well plates at 0.6 × 10⁶ cells per well in DMEM containing 10% BGS, 1% penicillin–streptomycin, and 50 ng/mL M-CSF. The next day, BMDM medium was removed and wells were washed with 1× PBS. TGFβ-stimulated fibroblast conditioned medium was then transferred onto BMDMs for 24 h, and cells were collected for Plasmidsaurus bulk RNA sequencing or qPCR analysis.

### RNA extraction, RT-PCR, and real-time PCR analysis

Total RNA was extracted using the Direct-Zol RNA kit (Zymo Research) according to manufacturer instructions. 500 ng of RNA was converted to cDNA using iScript cDNA synthesis kit (Bio-Rad). For Taqman real-time PCR analysis, 1/50 cDNA was used for quantitative PCR with Taqman Universal PCR master mix (Life Technologies) and the QuantStudio 5 Real-Time PCR System (ThermoFisher Scientific). For SYBR Green real-time PCR analysis, 1/10 of the cDNA was used for quantitative PCR with SYBR Green Supermix (Bio-Rad) on a CFX384 Real-Time PCR Detection System (Bio-Rad). For The Taqman probes and SYBR primers are outlined in the relevant “Tables” section.

### Cross-organ fibroblast atlas re-analysis

Processed Seurat objects containing steady-state and perturbed fibroblast single-cell RNA-seq data were downloaded from the Buechler et al. cross-organ fibroblast atlas (fibroxplorer.com; 220,179 fibroblasts, 16 tissues, 28 disease conditions, 13 transcriptionally defined cell states)^20^. The two objects were merged, log-normalized (NormalizeData, scale.factor = 10,000), and ALPINE gene program activity was scored per cell using Seurat’s AddModuleScore (top 25 genes per program by NMF loading weight; nbin = 24, ctrl = 100, seed = 42), yielding 15 per-cell scores. For injury kinetics, mean module scores were computed per program × organ × condition across three fibrosis models spanning acute, chronic, and resolving trajectories: lung (healthy → bleomycin day 11 → resolution day 21), liver (healthy → CCl₄ acute → CCl₄ chronic), and skin (healthy → excisional wound). No batch correction or weighting was applied; effect sizes are reported as Δ = mean(first disease timepoint) − mean(healthy).

### Definition of the Fibrosis Gene Signature for dCas12 experiment

We defined a fibrosis gene signature comprising genes that were dynamically modulated in a murine model of heart failure. Specifically, we used a transverse aortic constriction (TAC) model with intermittent exposure to the potent anti-fibrotic BET bromodomain inhibitor JQ1 (PMID: 34163071). In this model, profibrotic gene transcription is activated by TAC-induced pressure overload, strongly repressed during JQ1 treatment, and re-upregulated upon JQ1 withdrawal, demonstrating that these genes are dynamically and reversibly engaged during the fibrotic response. This yielded 169 dynamically modulated genes. From the 169 TAC-JQ1 dynamic genes, we retained only those that were also expressed in human cardiac fibroblasts in chronic heart failure (HF) or acute myocardial infarction (AMI), reducing the signature to 130 genes. Having established which enhancers are profibrotic in nature in our CROP-seq study (Figure 2A,B; 103 enhancers), we further retained only those signature genes whose expression was modulated by at least one of these profibrotic enhancers. This yielded a final fibrosis gene signature of 110 genes. Using the CROP-seq data, we determined how many enhancers must be repressed to suppress increasing fractions of this 110-gene fibrosis signature. Enhancer combinations were prioritized based on enhancers assigned to ALPINE categories previously identified as key regulators of the profibrotic transcriptional state: ECM & Matrix Maturation (P9), Contractile Myofibroblast (P5, P6, P10), Myofibroblast Proliferation (P13, P14) and Migration & Paracrine (P7). In the saturation analysis, enhancers were iteratively added in order of the number of uniquely affected fibrosis signature genes: E1 (48 genes), E4 (15 genes), E2 (10 genes), E6 (6 genes), E5 (5 genes), E3 (3 genes). This analysis revealed a saturation curve in which repression of just 6 enhancers was sufficient to silence 87 of the 110 fibrosis genes, with diminishing returns for additional enhancers. These 6 enhancers were subsequently targeted simultaneously using the dCas12a-KRAB multiplexed CRISPRi system. The six enhancers targeted by the array are referred to as E1–E6: E1 (chr18:33,667,058–33,668,110; P7 Migratory-Paracrine), E2 (chr16:93,246,720–93,247,220; P9 ECM Deposition), E3 (chr18:35,889,526– 35,890,026; P6 Tagln+ MyoFB), E4 (chr13:56,599,750–56,602,086; P10 Myl9+ MyoFB), E5 (chr1:92,378,979–92,379,704; P13 Postn+ Proliferation), and E6 (chr5:53,542,987–53,543,961; P4 MyoFB Mitosis).

### Generation of dHyperLbCas12a-KRAB (dCas12a) Fibroblast Cell Line

The dCas12a fibroblast cell line was generated using a previously established immortalized mouse cardiac fibroblast cell line^5^. The immortalized mouse cardiac fibroblast line was transduced with crude lentiviral medium containing the SM266_EF1A-dHyperLbCas12a-KRAB-T2A-BlastR plasmid (gift from Dr. Timothy Reddy, Addgene #244205) supplemented with 8 µg /mL of polybrene (MilliporeSigma TR-1003-G). Lentiviral medium was removed 24h post transduction and replaced with normal culturing medium. Cells with lentivirus integration were then selected for using 5 ug/mL of Blasticidin S HCl (ThermoFisher Scientific J67216) and maintained at 2.5 µg /mL. Blasticidin resistant cells were then single-cell sorted (BD Aria Fusion) into 96-well plates to generate a dCas12a CRISPRi clone.

### Generation of the gRNA-Array dCas12a CRISPRi Line

crRNAs targeting the six CROP-seq nominated enhancers were designed using CRISPick, selecting spacers adjacent to a Cas12a TTTV PAM. Two crRNAs were selected per enhancer, yielding a 12-crRNA array in which each spacer was separated by the dHyperLbCas12a (Lb) direct repeat (DR). The array was synthesized and cloned into the SM425 lentiviral pre-crRNA expression vector backbone (gift from Dr. Timothy Reddy, Addgene #244225) by GeneWiz (Azenta) and sequence-verified by Quintara whole-plasmid Nanopore sequencing. To generate the gRNA-array line, the stable dHyperLbCas12a-KRAB effector line was transduced with 2 mL of undiluted crude lentivirus per well of a six-well plate. The gRNA selected were outlined in the relevant “Tables” section. The array construct carries an mCherry marker driven by a CAG promoter, allowing transduced cells with the gRNA-array to be enriched by fluorescence-activated cell sorting (FACS), gating on the brighter ∼80% of the mCherry-positive population to exclude low-expressing cells.

### Lentivirus Production for dCas12a CRISPRi Experiments

Lentivirus was produced in HEK293T cells. 3 × 10⁶ HEK293T cells were seeded per 10-cm dish in DMEM (Gibco 10566-019) supplemented with 10% FBS, 1 mM sodium pyruvate, 1× non-essential amino acids, and 10 U/mL penicillin/streptomycin. Cells were allowed to recover for 24h before the medium was replaced with 10 mL fresh medium, and cells were transfected with 5 µg of the lentiviral expression plasmid, 2.5 µg pMD2.G, and 2.5 µg psPAX2 using 59 µL FuGENE HD (Promega E2312) in a total of 1,179.5 µL Opti-MEM (Gibco 31985062). FuGENE HD was first added to the Opti-MEM, mixed, and incubated for 5 min at room temperature; the plasmid mixture was then added, mixed, and incubated for a further 15 min before being added dropwise to the cells. Viral supernatant was collected 48h post-transfection and clarified through a 0.45 µm syringe filter. The same procedure was used to produce both the dCas12a-effector and pre-crRNA array lentiviruses.

### Network analysis for dCas12

Differential expression was computed with limma (moderated t-test, batch-aware where applicable) with Benjamini–Hochberg adjustment, and genes called at FDR < 0.05 and |log₂FC| > 0.5; differential ATAC peaks (FDR < 0.05, |log₂FC| > 1) were assigned to genes using ENCODE-rE2G enhancer–gene predictions for TGFβ-treated cardiac fibroblasts, each peak to every gene in its prediction set. Enrichment of lost enhancers among repressed genes, and overlap between the enhancer and knockdown gene sets, were tested by two-sided Fisher’s exact test against all measured genes; enrichment of the 6Enh-repressed set within each knockdown ranking used a weighted running-sum statistic with 2,000 gene-label permutations. Mouse and human genes were matched by uppercase symbol as a 1:1 orthologue proxy. Gene sets were taken unmodified from MSigDB v7.0 (Hallmark and GO biological process), with terms selected by name before any result was inspected and tested by two-sided Mann–Whitney of set-gene log₂ fold changes against all other measured genes with BH correction across all set-by-perturbation comparisons; effect size is reported as the mean log₂ fold change, since the largest sets reach significance on negligible shifts. Transcription factors were defined by the pySCENIC mouse MGI annotation, and transcription factor → program edges are literature-based rather than inferred. Analyses used Python 3.12 (pandas, numpy, scipy) with matplotlib figures.

### *Meox1^Flox^* Genotyping Primers

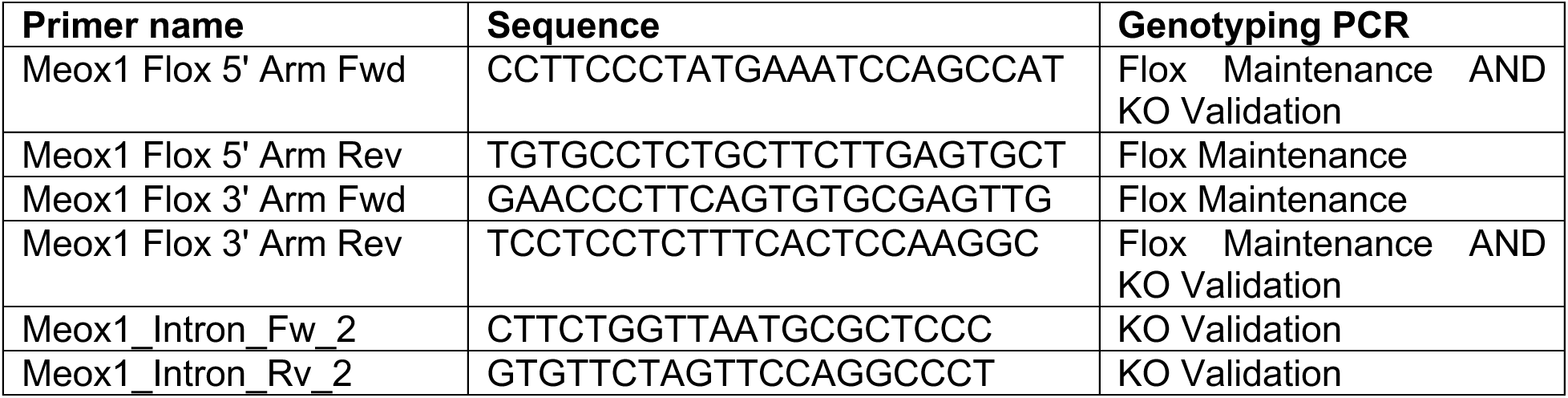

### CROP-seq Vector Modification

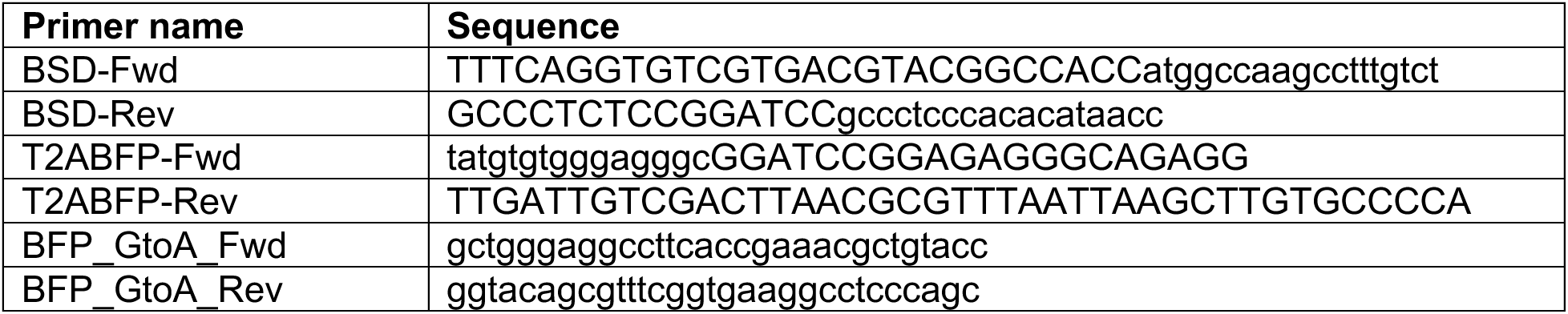

### sgRNA Amplicon Library Generation

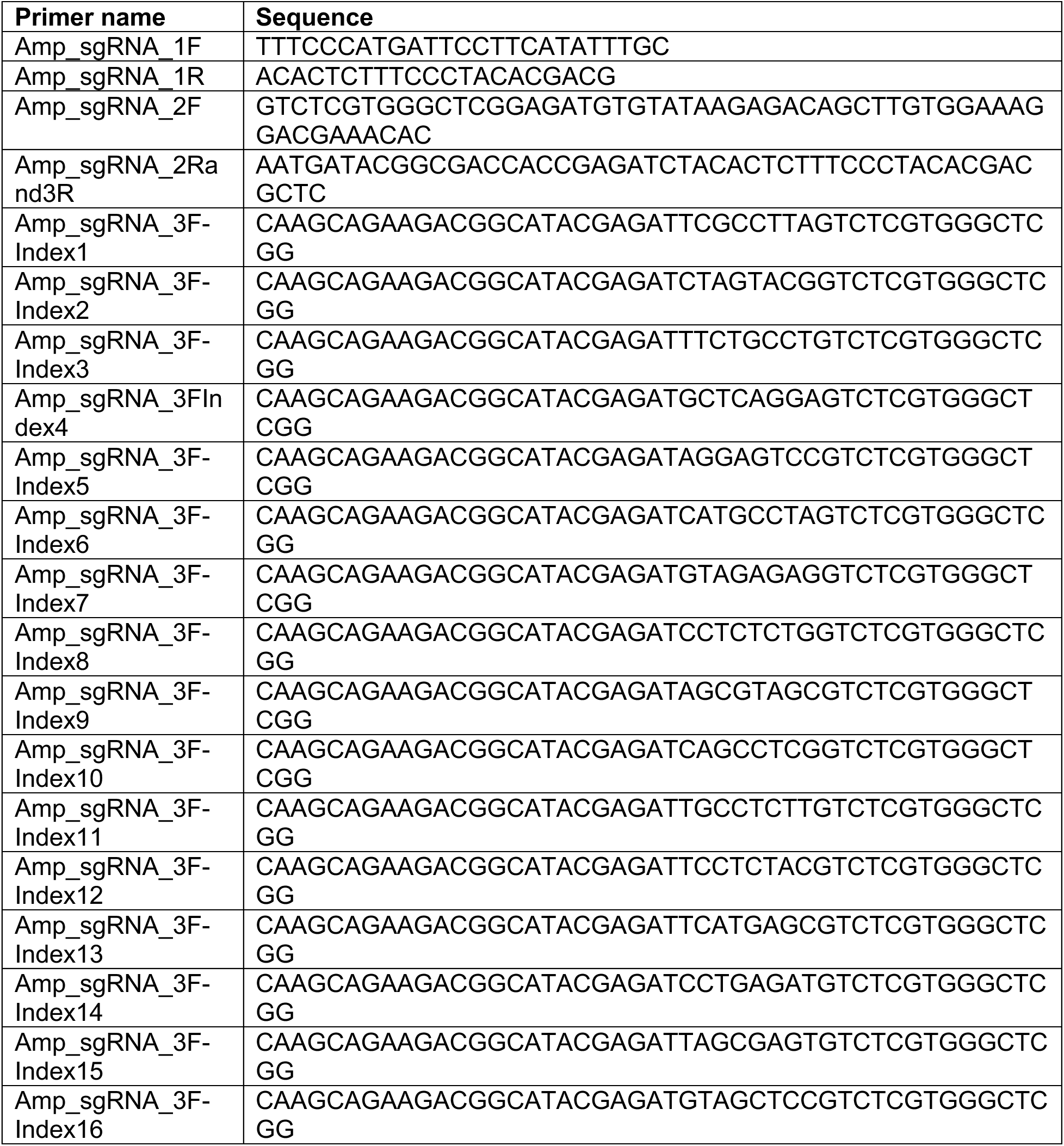

### Generation of the gRNA-Array dCas12a CRISPRi Line

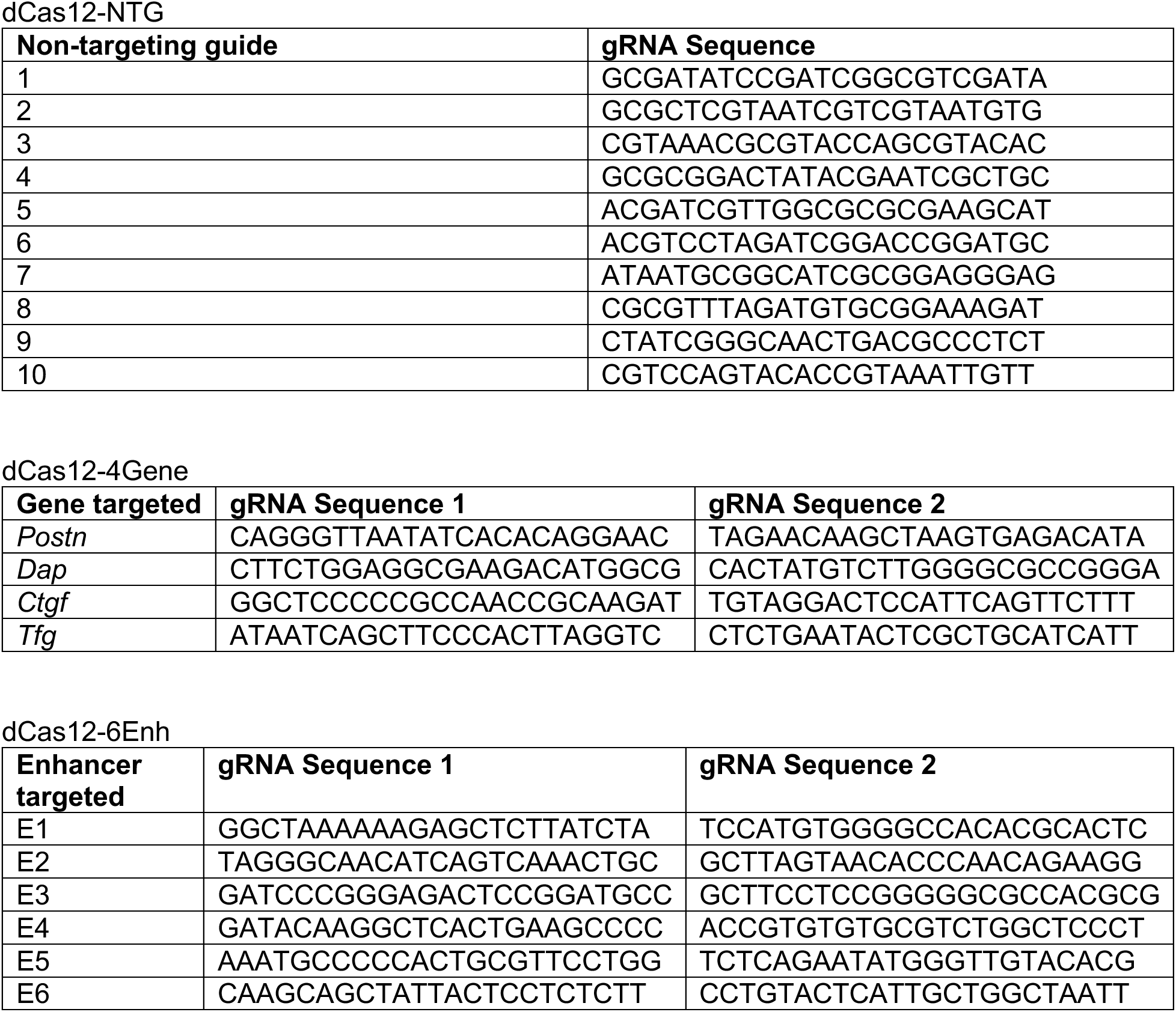

### siRNAs used on human primary cardiac fibroblasts

Dharmacon ON-TARGETplus SMARTpool siRNAs

siRNAs used on human primary cardiac fibroblasts Dharmacon ON-TARGETplus SMARTpool siRNAs
| Gene | Lot# |
| --- | --- |
| *ADAM12* | L-005118-00 |
| *EDN1* | L-016692-00 |
| *LRRC32* | L-012100-01 |
| *SERPINE1* | L-019376-01 |
| *COL5A1* | L-011015-00 |
| *BGN* | L-021493-00 |
| *PSD3* | L-014030-01 |
| *SOCS5* | L-017374-00 |
| *MICAL2* | L-010189-00 |
| *AEBP1* | L-009694-00 |
| *TGFBI* | L-019370-00 |
| *GPC1* | L-004303-02 |
| *RBPJ* | L-007772-00 |

### CRISPRi knockdown of transcription factors (gRNAs used)

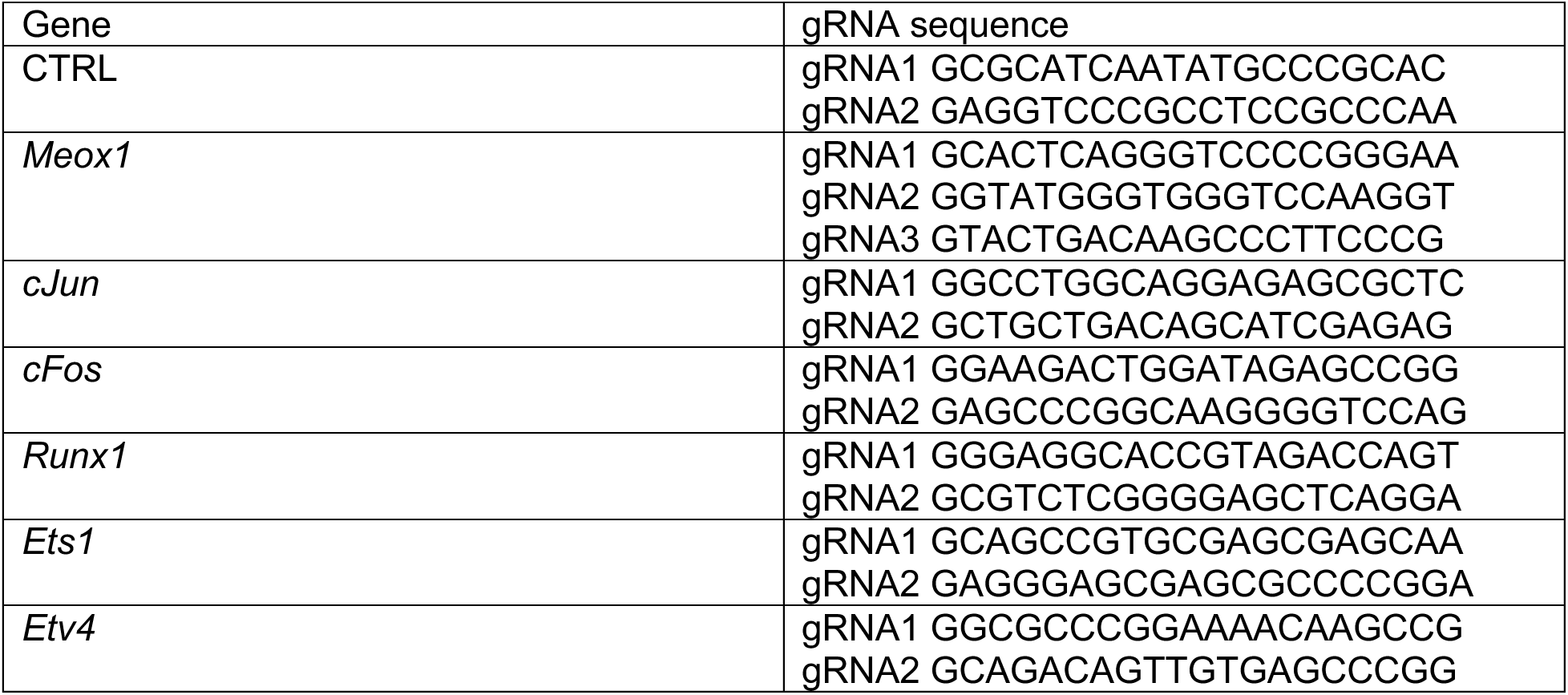

### Human Sybr primers for RT-qPCR

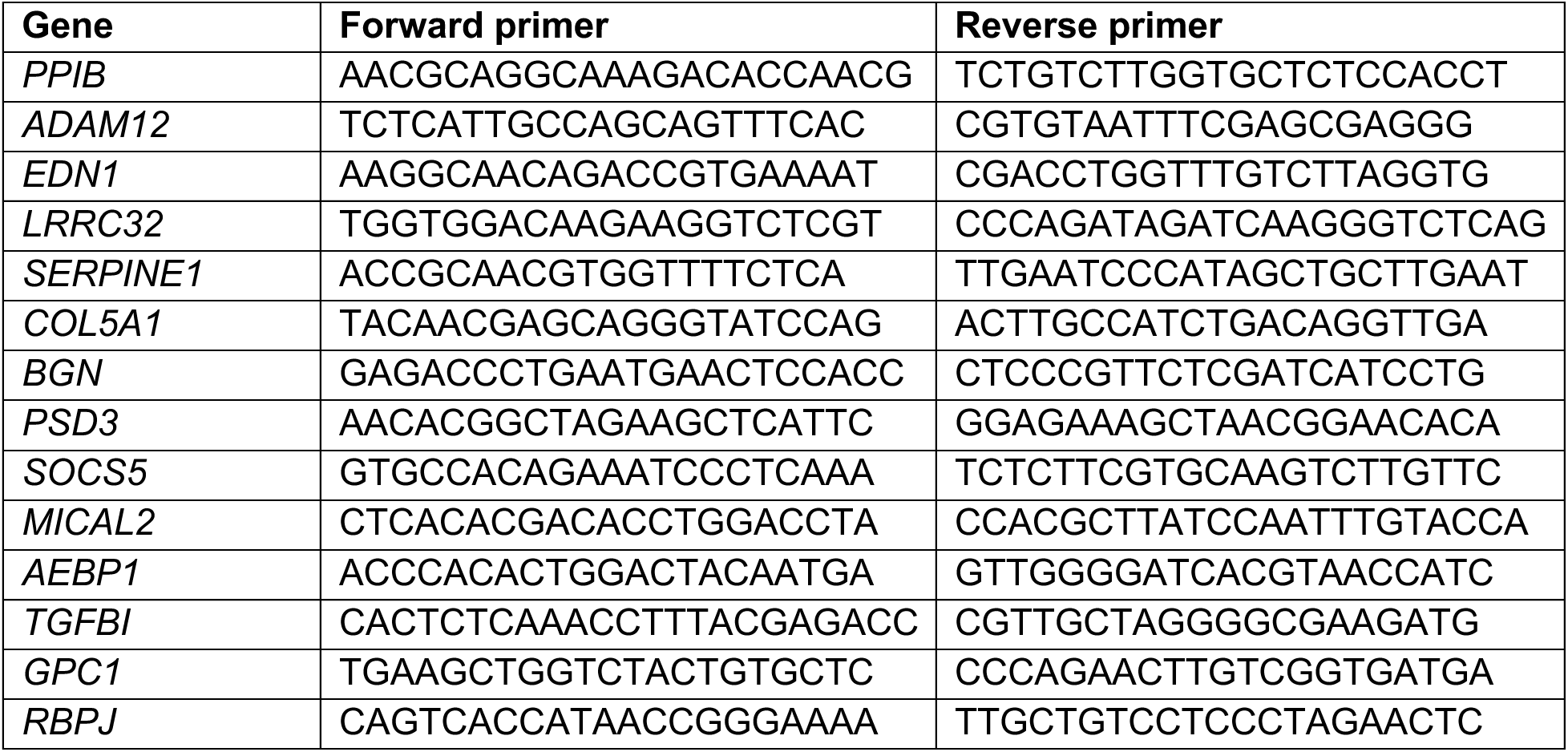

### Mouse Sybr primers for RT-qPCR

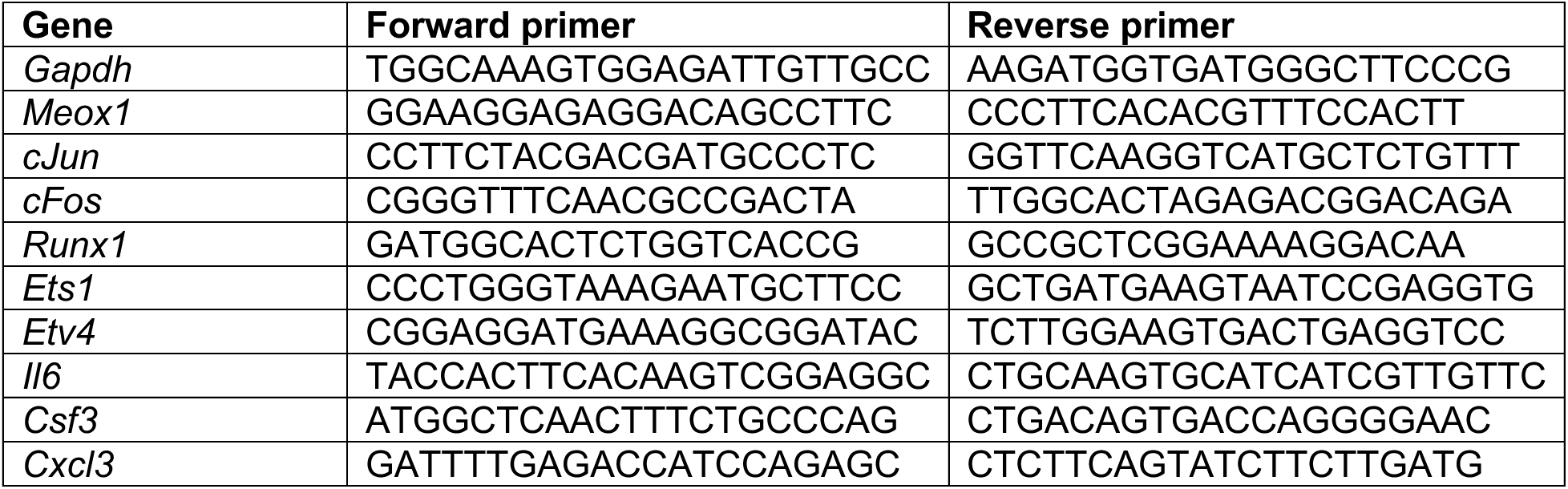

